# Metagenomics and Metatranscriptomics Suggest Pathways of 3-Chloroaniline Degradation in Wastewater Reactors

**DOI:** 10.1101/2021.05.02.442374

**Authors:** Hari Seshan, Ezequiel Santillan, Florentin Constancias, Uma Shankari Chandra Segaran, Rohan B. H. Williams, Stefan Wuertz

## Abstract

Biological wastewater treatment systems are often affected by major shifts in influent quality, including the input of various toxic chemicals. Yet the mechanisms underlying adaptation of activated sludge process performance when challenged with a sustained toxin input are rarely studied in a controlled and replicated experimental setting. Three replicate bench-scale bioreactors were subjected to a chemical disturbance in the form of 3-chloroaniline (3-CA) over 132 days, after an acclimation period of 58 days, while three control reactors received no 3-CA input. Nitrification was initially affected by 3-CA but the microbial communities in all three treatment reactors adapted to biologically degrade 3-CA within three weeks of the experiment, resulting in partial nitrification recovery. Combining process and microbial community data from amplicon sequencing with potential functions gleaned from assembled metagenomics and metatranscriptomics data, two putative degradation pathways for 3-CA were identified. The first pathway proceeds via a phenol monooxygenase followed by ortho-cleavage of the aromatic ring, and the second one involves a benzoate dioxygenase and subsequent meta-cleavage of the aromatic ring. The genera *Gemmatimonas*, *OLB8*, and *Taibaiella* correlated significantly with 3-CA degradation. Metagenome-assembled genome data also showed the genus *OLB8* to be differentially enriched in treatment reactors, making it a strong candidate as 3-CA degrader. Using replicated reactors, this study has demonstrated the impact of a sustained stress on the activated sludge community and processes carried out by its members, followed by process recovery. By a combination of techniques, we showed that microbial communities can develop degradative capacity following a sustained xenobiotic input, and that targeted culture-independent approaches can suggest plausible mechanisms for 3-CA degradation and identify the taxa potentially contributing to it.

## 1 Introduction

The activated sludge process in biological wastewater treatment involves a complex microbial community performing several metabolic functions under controlled conditions (Briones and Raskin, 2003). Major shifts in process performance can occur when a disturbance is imposed on a community (Allison and Martiny, 2008) and can impact a wastewater treatment plant’s treatment capability over the long-term (Vuono et al., 2015). Such disturbances may be operational (Thompson, 2006; Valipour et al. 2014, Santillan et al., 2020), physical (Hwang et al., 2006; Vargas et al., 2013), or chemical (Kelly et al., 2004; Falk and Wuertz, 2010). While short-term (pulse) disturbances have been studied extensively (Boon et al., 2000; Henriques et al., 2007; Ma et al., 2020), long-term (sustained) disturbances have not. Sustained disturbances provide an opportunity for processes to adapt and recover (Johnston et al., 2019), and studying them may provide insights into means to buffer systems against disruptions of treatment processes (Briones and Raskin, 2003).

The compound 3-chloroaniline (3-CA) is a by-product of the production of herbicides and dyes (Ficara and Rozzi, 2001). It disrupts nitrification by inhibiting ammonia oxidizing bacteria (AOB) (McCarty, 1999; Falk and Wuertz, 2010). Its impact on wastewater treatment has been assessed using pulse inputs ranging from 40 mg/L 3-CA (Ma et al., 2020) to 250 mg/L 3-CA (Boon et al., 2000), continuous inputs up to 120 mg/L 3-CA (Falk and Wuertz, 2010; Zhu et al., 2008), and recurring pulse inputs of 70 mg/L at varying frequencies (Santillan et al., 2019). Biological 3-CA degradation involves its conversion to a chlorocatechol, followed by cleavage of the aromatic ring and channelling into TCA cycle intermediates (Reineke, 1998; Pimviriyakul et al., 2019). Some of these studies reported 3-CA degradation (Zhu et al., 2008; Santillan et al., 2019), but none confirmed the pathway involved or identified the microbial community members contributing to degradation.

Metagenomic and metatranscriptomic analyses allow the examination of gene abundance and expression, respectively, in complex microbial communities like activated sludge. Transcriptomics has been used to elucidate degradation pathways in the context of aromatics degradation in pure culture experiments (Tam et al., 2006; Yan and Wu, 2017; Cauduro et al., 2020; Ma et al., 2020b). Long-term metatranscriptomics sampling has also shown differential abundance and expression of genes related to aromatic-compound degradation in activated sludge bioreactors (Sato et al., 2019; Sun et al., 2019; Yu et al., 2019). However, it is rare to combine these methods with amplicon sequencing to determine the taxa involved in degradation and outline possible catabolic pathways. Furthermore, the expression of relevant genes in a bioprocess can potentially be followed using metatranscriptomics but is rarely exploited in the context of process engineering.

This study investigated the effect of a sustained xenobiotic disturbance in the form of 3-CA addition for 132 days on activated sludge wastewater treatment, focusing on process impacts and 3-CA degradation mechanisms. Of six bench-scale bioreactors that had been acclimated with activated sludge from a wastewater treatment plant, three received sustained 3-CA input and three were maintained as controls, with no 3-CA input. The objectives of this study were to: (a) examine, through intensive nutrient cycling profiles, how the emergence of 3-CA degradation impacted process recovery and the nitrifier microbial community; (b) identify, via amplicon sequencing and metagenomics, the taxa most likely responsible for 3-CA degradation; and (c) identify, via metagenomics and metatranscriptomics, the most likely degradation pathways involved in 3-CA degradation.

## 2 Materials and methods

### 2.1 Experimental design

Six sequencing batch bioreactors were run in parallel, fed on synthetic wastewater, for a 58-day preliminary acclimation period. On day 0 (D0), the inoculum was taken from aeration tanks of a full-scale water reclamation plant in Singapore, mixed and distributed to all six reactors (2 L each). Each reactor additionally received 2 L of synthetic wastewater for a working volume of 4 L. The reactors were run identically on nonstop 12-h cycles during acclimation.

The stages of each 12-h cycle were as follows: 20 min feeding, 180 min anoxic mixing, 440 min aeration (dissolved oxygen, DO, maintained at 1-2 mg/L using a feedback loop with aeration beginning at 1 L air/min when probes measured DO < 1 mg/L, and stopping when DO > 2 mg/L), 50 min settling and 30 min effluent (supernatant) discharge. 2 L of effluent were discharged at the end of every cycle and replaced with 2 L of synthetic wastewater at the beginning of the next 12-h cycle, resulting in a hydraulic retention time of 24 h. The mixed liquor temperature was maintained at 30□C using water jackets around the reactors and a re-circulating water heater. Solids were removed regularly from mixed liquor to maintain an SRT of about 30 d.

The synthetic wastewater fed to all six reactors in the feed phase of each cycle during the acclimation period was adapted from Hesselmann et al. (1999) and is detailed in Supplementary Information (SI). When diluted in reactor mixed liquor, it provided 500 mg/L COD and 70 mg/L Total Nitrogen.

On D59, the 3-CA experiment was started and continued for 132 days. At the start of this experiment, three reactors were randomly assigned to the treatment group (with 3-CA added to their feed regularly as described below) and the other three were assigned to the control group (no 3-CA addition). The reactors were originally numbered 1 through 8, with reactors 6 and 8 being removed from service during acclimation. Reactors 1, 3 and 7 were then randomly selected to serve as treatment reactors, while 2, 4 and 5 served as controls. Treatment reactors 1, 3 and 7 were subsequently redesignated T1, T2 and T3 respectively, and control reactors 2, 4 and 5 were redesignated C1, C2 and C3 respectively. This redesignation is used in this paper. Cycle conditions and other parameters were identical to acclimation phase conditions, except that treatment reactors were fed differently: the organic constituents in treatment reactors were reduced by 20%, to a COD of 400 mg/L. The remaining 20% of COD was as 3-CA, resulting in a mixed liquor 3-CA concentration of 70 mg/L and a total COD concentration (when diluted in mixed liquor) of 500 mg/L. The three control reactors continued to receive the same 3-CA-free medium used during acclimation. The 3-CA experiment lasted 132 days, or roughly 4 SRTs. On D99, a pulse 3-CA load was inadvertently added to control reactor C3 at the same concentration as the treatment reactors, as detailed in SI. The majority of results from the time points shown in main figures are either prior to this error, or after its impacts had dissipated. In other words, data from reactor C3 from after this error were captured after process performance had recovered to its pre-error levels.

### 2.2 Mixed liquor sampling for molecular analyses

To assess the changes in community composition, community function, and treatment performance over the long term, samples of mixed liquor and effluent were taken from all six reactors at five distinct time points during the experiment; specifically, on D0 (inoculation of all six reactors), D56 (shortly before 3-CA input began), D63 (shortly after 3-CA input began), D112, D164 and D176. The last three sampling dates represent periods where 3-CA degradation had fully developed in treatment reactors. Mixed liquor was collected and processed as described in Santillan et al. (2021). Mixed liquor was also sampled hourly during a single cycle study (see Section 2.3) on D171 for RNA extraction and sequencing as described in Section 2.8.

### 2.3 Sampling for full-cycle process performance assessment

To investigate nutrient cycling across all six bioreactors, intensive sampling was carried out during 12-h cycles roughly every two weeks during the experiment (termed cycle studies). Ten cycle studies were conducted throughout the 3-CA experiment, on D59 (first day of 3-CA addition to treatment reactors), D80, D95, D110, D124, D129, D136, D158, D171 and D192. Results from five representative cycle studies (D59, D95, D124, D158 and D192) are presented in main figures, while all cycle study results are in SI (Figures S1 – S5).

During a cycle study, samples were taken from all reactors at several time points. The first cycle study sample was usually of of effluent (supernatant) from the end of the previous cycle, taken at the end of settling, 30 min before the cycle under study commenced. Mixed liquor samples were then taken hourly during the anoxic phase, starting 30 min after the cycle started, up to the anoxic-aerobic interface. During the aerobic phase, mixed liquor was sampled every 90 min (D59 to D129) or every 30 min (D136 to D192), until the beginning of the settling phase. Then a final effluent sample was taken at the end of the settling phase. All samples were centrifuged at 8000*g* for 3 min, and the supernatant filtered through a 0.2-μm filter for water quality analyses as described in Section 2.4.

Note that mixed liquor for molecular analysis was not sampled on the same days as for cycle studies. This is because cycle studies were used for short-term assessments, whereas the molecular (DNA) methods were primarily targeted at monitoring long-term community changes. Note however that the dates for long-term sampling are close to cycle study dates.

### 2.4 Water quality parameters

Water quality parameters were measured as described by Santillan et al. (2021) and detailed in SI.

### 2.5 DNA extraction

Genomic DNA was extracted and processed as described by Santillan et al. (2021) and detailed in SI.

### 2.6 16S rRNA gene amplicon sequencing and bioinformatics

Extracted DNA from D0, D56, D63, D112 and D164 were amplified for the 16S rRNA gene and the amplicons were sequenced using the Illumina MiSeq (v.3) platform. The amplification, sequencing and accompanying bioinformatics are described by Santillan et al. (2021) and detailed in the SI. To allow for consistent scaling of relative ASV abundance, the read counts of ASVs identified in each sample were normalized to the minimum total read count seen in any given sample. The relative abundances of ASVs identified in the normalized matrix were used to identify correlations with 3-CA degradation as described in Section 2.9.1. Whole microbial community-level analysis of these data is available elsewhere (Santillan et al., 2021).

### 2.7 Metagenomics sequencing and bioinformatics

Mixed liquor from all six reactors on D176 of the experiment was used for whole community shotgun metagenome sequencing. Genomic DNA was extracted as described in Section 2.5 and sequenced at SCELSE on an Illumina HiSeq platform (Illumina, Inc., San Diego, California, USA). Prior to library preparation, the quality of the DNA samples was assessed on a Bioanalyzer 2100 using a DNA 12000 Chip (Agilent Technologies, Santa Clara, California, USA). Library preparation was performed following Illumina’s TruSeq DNA Nano Sample Preparation protocol, starting with 100 ng of genomic DNA and following sizing instructions for 350bp inserts. During adapter ligation, each library was tagged with one of Illumina’s TruSeq LT DNA barcodes. Library quantification was performed using Invitrogen’s Picogreen assay and average library size was determined by running libraries on an Agilent Bioanalyzer DNA 7500 chip. Library concentrations were normalized to 4 nM and validated by qPCR on an Applied Biosystems ViiA-7 real-time thermocycler using qPCR primers recommended in Illumina’s qPCR protocol, and Illumina’s PhiX control library as standard. Libraries were then pooled and sequenced across two lanes of an Illumina HiSeq 2500 rapid sequencing run at a read-length of 150-bp paired-end.

In total, around 634 million paired-end reads were generated, averaging 105 ± 3 million paired-end reads per sample. Functional gene identification and quantification was performed using the SqueezeMeta pipeline (Tamames and Puente-Sánchez, 2019). Read pairs were co-assembled using Megahit (Li et al., 2015), generating 2.1 million contigs representing 2,649 million nucleotides. Open reading frames (ORFs) were then predicted from contigs using Prodigal (Hyatt et al., 2010), from which 12 million ORFs were identified, averaging 2 ± 0.1 million ORFs per sample. Functional annotation was performed using DIAMOND (Buchfink et al., 2015) against the KEGG database (Kanehisa et al., 2016). Read mapping against contigs was then performed using Bowtie2 (Langmead and Salzberg, 2012) to quantify the abundance of genes among the different samples, and transformed to transcripts per million (TPM) values (Puente-Sánchez et al., 2020). To identify genes involved in 3-CA degradation, the analysis focused on the following KEGG metabolic pathways from the xenobiotics biodegradation and metabolism category: chlorocyclohexane and chlorobenzene degradation (KO: 00361), benzoate degradation (KO: 00362), polycyclic aromatic hydrocarbon degradation (KO: 00624), and aminobenzoate degradation (KO: 00627). 170 genes were recovered in total from these pathways.

Metagenome-assembled genome (MAG) binning was performed using MaxBin2 (Wu et al., 2016), Metabat2 (Kang et al., 2019) and DAS Tool (Sieber et al., 2018) using the above-mentioned co-assembly as well as sample read mapping quantification. MAG genome quality statistics were computed using CheckM (Parks et al., 2015). Overall, 60 MAGs were reconstructed from the dataset at <10% contamination and >80% completeness, the latter exceeding the >50% completeness required for medium-quality MAGs (Bowers et al., 2017). Genome level taxonomy was assigned using GTDB (Chaumeil et al., 2020, release 06-RS202) and functional potential was further estimated using anvio (Eren et al., 2015) against KEGG Orthologs (KOs) (using anvi-setup-kegg-kofams, anvi-run-kegg-kofams, default parameters). Quantification of MAG relative abundance was performed using mean coverage across samples. A summary of the statistics from the metagenomic analyses is available as part of Supplementary Files (SF).

### 2.8 RNA extraction, sequencing and transcriptomics

To determine the changes in gene expression profiles during a single 12-hour cycle, RNA extraction and subsequent sequencing were performed on each sample taken from a single cycle study on D171. Total RNA extraction methods have been described in Law et al (2016). RNA extractions were performed with Soil/ Fecal RNA Isolation Kit (Zymo Research, Irvine, CA, USA; Catalog# R2040) with some protocol modifications (Law et al, 2016). RNA purity was determined by Thermo Scientific Nanodrop 2000 by calculating an A260/A280 and A260/A230 to screen for contaminants in purified RNA. Extracted RNA concentration was determined by a Qubit 2.0 fluorometer, and RNA integrity (RIN) was measured using the Agilent 2200 Tapestation system, prior to library preparation and sequencing on an Illumina 2500 using a 101□bp paired-end run (Law et al, 2016). Reads of non-ribosomal origin were mapped to predicted genes in the metagenome assembly, converted to reads per kilobase million RPKM) and aggregated using the same annotations described above.

### 2.9 Data Analysis

#### 2.9.1 Identification of ASVs associated with 3-CA degradation

To determine which ASVs from 16S rRNA amplicon sequencing were correlated with 3-CA degradation in treatment reactors, each ASV from treatment reactors from D64, D112 and D164 was compared to 3-CA degradation rates in treatment reactors on D59, D110 and D171, respectively. Same-day comparisons were not possible because 3-CA degradation data were not taken on the same days where mixed liquor (ML) samples were taken for amplicon sequencing. However, the variation in 3-CA degradation did not differ widely over time once 3-CA degradation had been established (Figure 1A; Figure S1), so these comparisons were still considered valid. Spearman’s rank order correlation coefficient was used between the three 3-CA removal percentages on the three days identified above, and the corresponding normalized ASV abundances in the three treatment reactors (n = 9: 3 time points × 3 reactors). To assess correlation significance, *t*-statistics were calculated from the correlation coefficient for each ASV based on 7 (n – 2) degrees of freedom, and *p*-values were calculated from these *t*-statistics. A correction for multiple comparisons was applied using the Benjamini-Hochberg procedure (McDonald, 2014), applying a 10% False Discovery Rate (FDR). The ASVs that remained significant (adjusted p-value < 0.05) after correction are discussed further in the next section. This analysis was implemented in Microsoft Excel.

**Figure 1.**
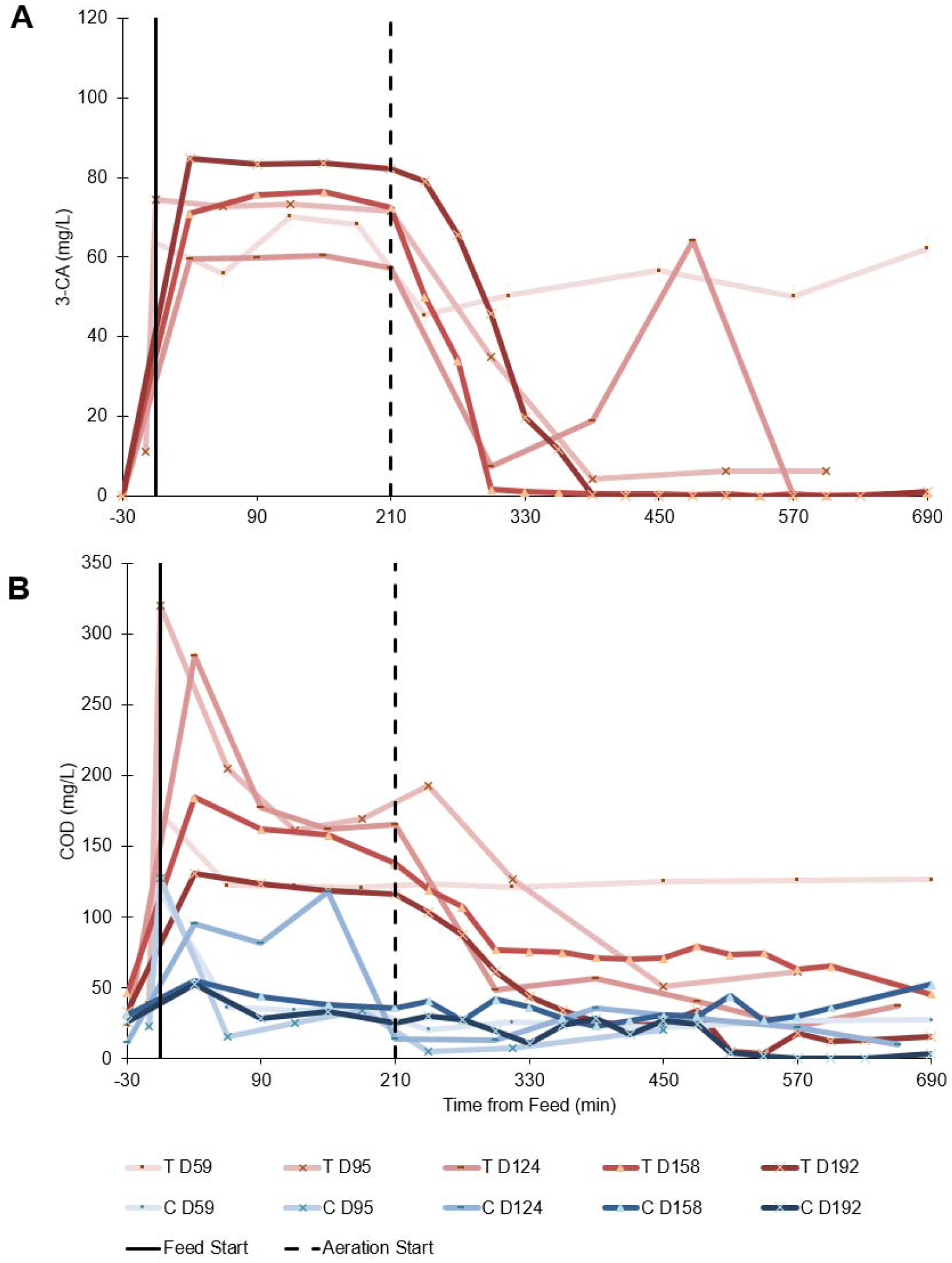
COD and 3-CA concentration profiles from five distinct days of the experiment showing single 12-h cycles for all six reactors. The points represent the mean concentrations of triplicate control and treatment reactors. The x-axes represent time from the commencement of feeding in min in each cycle. The dotted vertical line (at x = 210 min) separates the anoxic and aerobic phases. Aeration began at 210 min. The y-axes represent (**A**) 3-CA, mg/L and (**B**) COD, mg/L. Data series in shades of blue represent the mean of triplicate control reactors (no 3-CA) and data series in shades of red represent the mean of triplicate treatment reactors (3-CA added), with decreasing brightness over time. Note that the connecting lines between points are used to distinguish profiles visually and do not represent simulated or measured concentrations.

To examine these results visually, the mean normalized abundance of each taxon across all three treatment reactors on D63, D112 and D164 was plotted on a logarithmic scale against the respective 3-CA degradation correlation coefficient as calculated above (Figure 3). ASVs that fell toward the top right of this plot and correlated with relevant degradation genes as identified below were considered more likely to have participated in 3-CA degradation.

**Figure 2.**
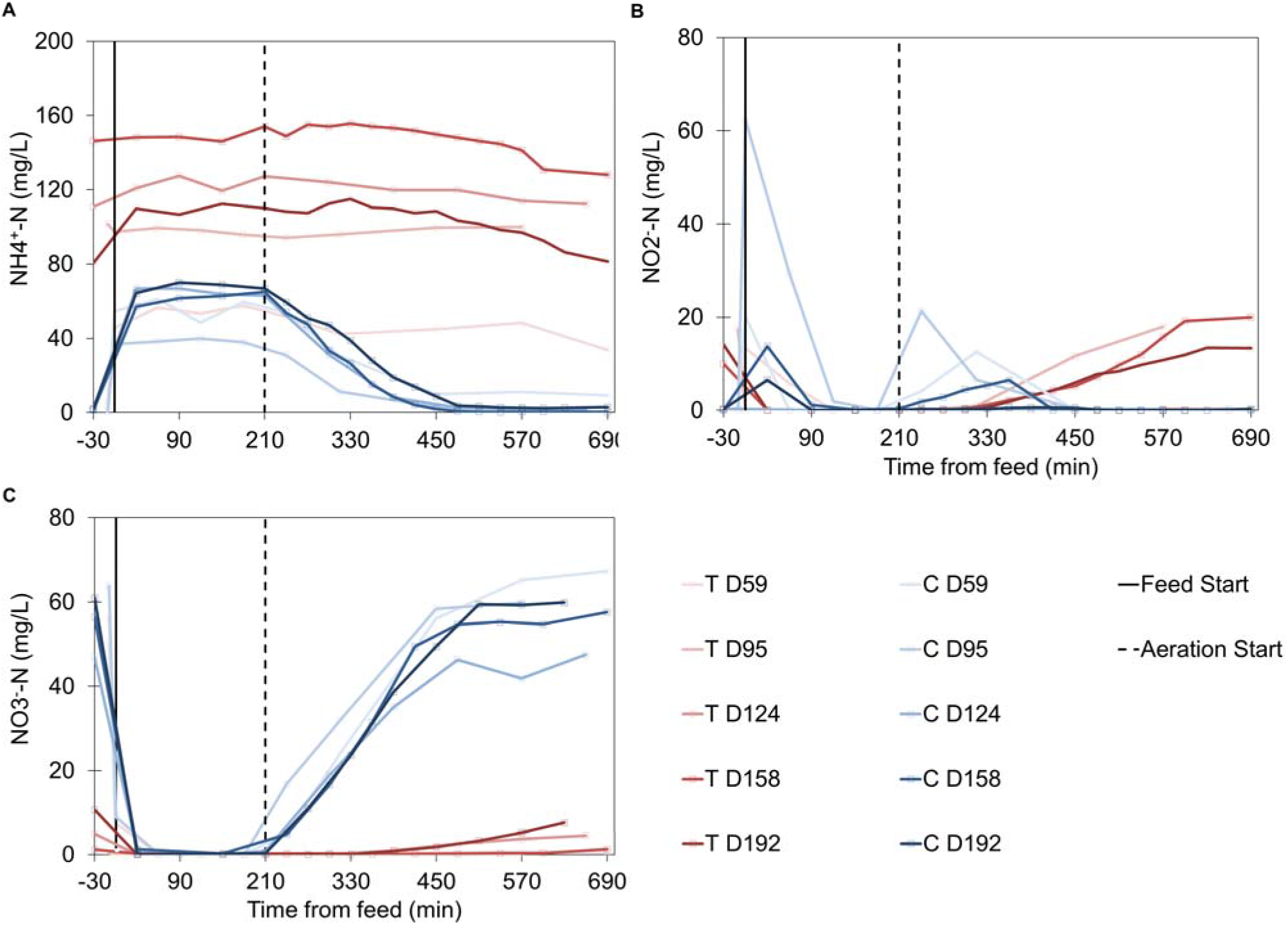
Ammonium, nitrite and nitrate concentration profiles from five distinct days of the experiment showing single 12-h cycles for all six reactors. The points represent the mean concentrations of triplicate control and treatment reactors. The x-axes represent time from the commencement of feeding in min in each cycle. The dotted vertical line (at x = 210 min) separates the anoxic and aerobic phases. Aeration began at 210 min. The y-axes represent (**A**) ammonium-nitrogen, mg/L NH_4_^+^-N, (**B**) nitrite nitrogen, NO_2_^−^-N mg/L and (**C**) nitrate nitrogen, mg/L NO_3_^−^-N. Data series in shades of blue represent the mean of triplicate control reactors (no 3-CA) and data series in shades of red represent the mean of triplicate treatment reactors (3-CA added), with decreasing brightness over time. Note that the connecting lines between points are used to distinguish profiles visually and do not represent simulated or measured concentrations.

**Figure 3.**
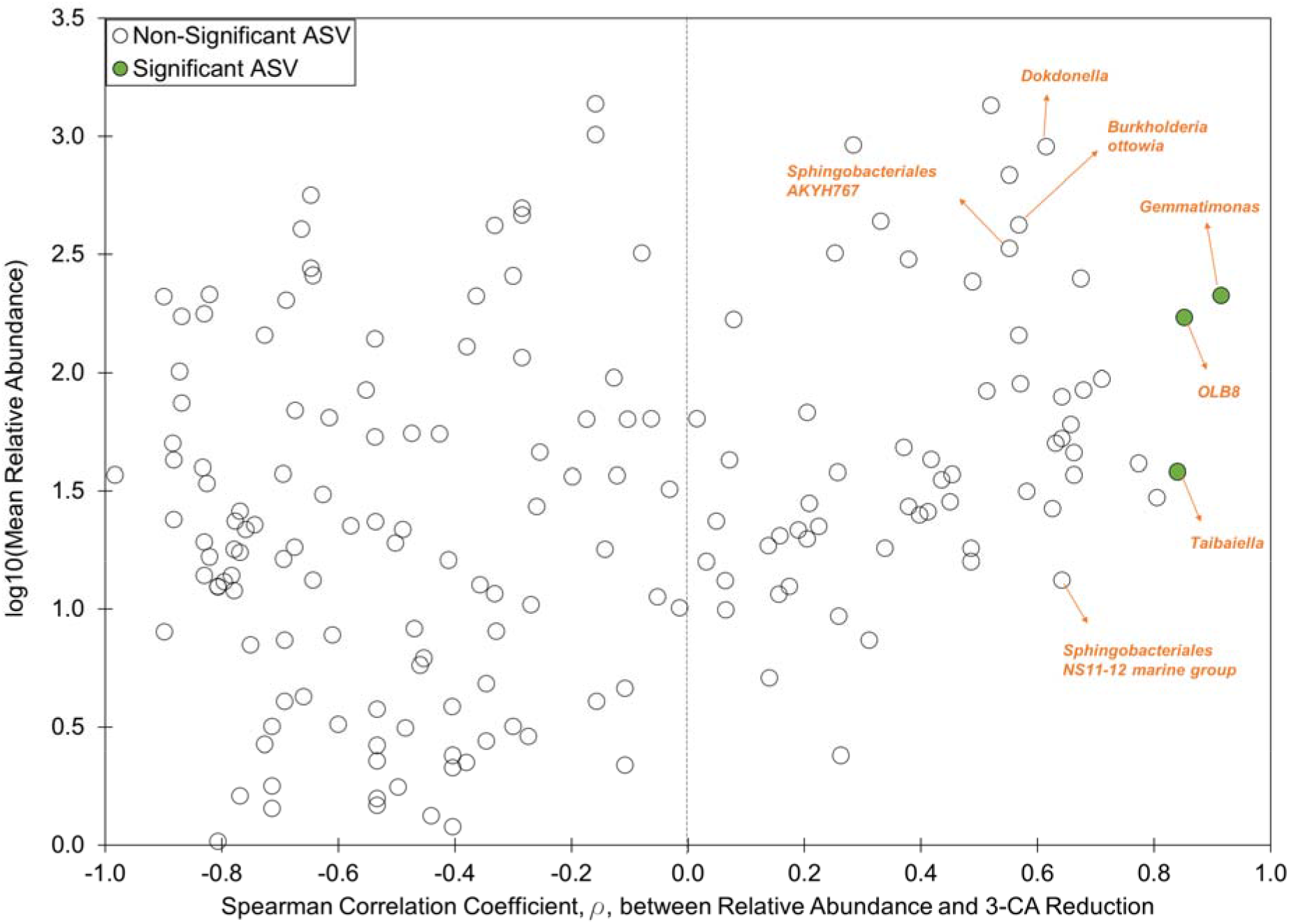
Relative abundances of the top 180 ASVs identified via 16S rRNA amplicon sequencing on D63, D112 and D163, plotted against the Spearman’s Rank Order Correlation Coefficient of their respective relative abundances to 3-CA reduction in treatment reactors on D59, D110 and D171, respectively (n = 9). ASVs with significant correlation to 3-CA degradation (filled in data points) after a 10% FDR adjustment for multiple comparisons are identified by name, as are other ASVs with high correlation that were considered relevant to 3-CA degradation.

#### 2.9.2 Enrichment of xenobiotics degradation genes

To determine which xenobiotics degradation genes identified from the metagenomics data set from D176 were differentially enriched, two methods were used in parallel. The first defined genes that were “differentially enriched” via calculating z-scores, while the second defined genes that were “significantly enriched” via general linear multivariate models (GLMMs). The overlap of these two sets produced a subset of genes that was examined more closely (Figure 4).

**Figure 4.**
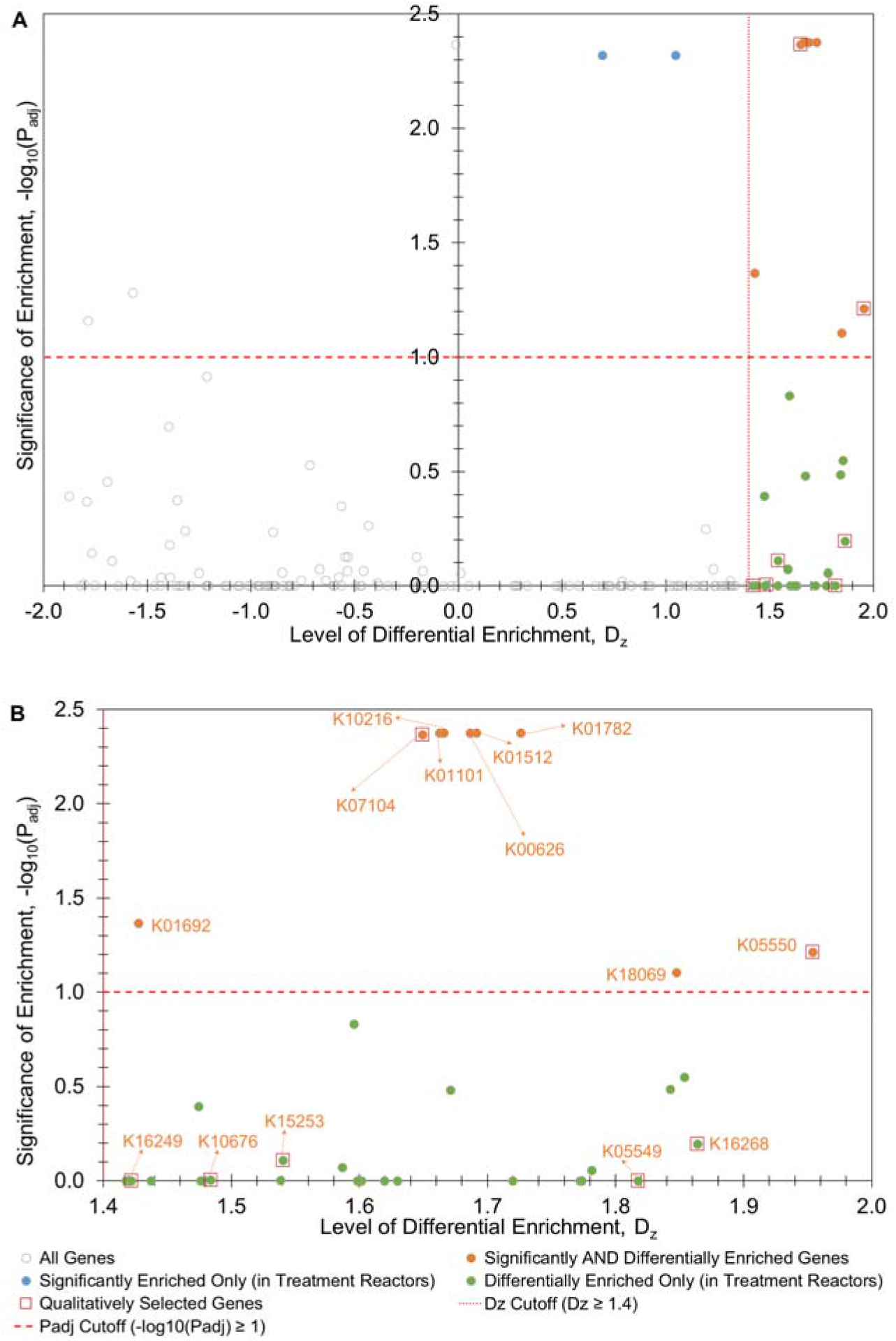
Levels of differential enrichment and significant enrichment in treatment reactors of (A) all 170 xenobiotics degradation genes identified in the metagenomics data set from D176, and (B) zoomed-in figure of the subset of these 170 genes that was differentially enriched (D_z_ ≥ 1.4). The negative base-10 logarithm of the adjusted p-value from the GLMM model (significance of enrichment) is plotted against levels of differential enrichment, D_z,_ for each gene. The adopted cut-off values for each method used (differential enrichment cut-off D_z_ ≥ 1.4; significant enrichment −log_10_(P_adj_) ≥ 0.1). Genes that are only differentially enriched in treatment reactors (23), only significantly enriched in treatment reactors (GLMM) (2), and genes enriched under both criteria (9), are coloured separately. Additionally, Figure 4B identifies genes that were both significantly and differentially enriched, as well as the 7 qualitatively selected genes by KEGG ortholog.

To evaluate differential enrichment, a z-score was first calculated from the TPM values for each gene in each reactor to determine the deviation from the average TPM. The difference in mean z-score between control and treatment reactors was then calculated, termed D_z_. Differential enrichment in treatment reactors was defined as D_z_ ≥ 1.4. This arbitrary cut-off balanced the selection of genes enriched by 3-CA presence and degradation, against the false elimination of genes that degraded 3-CA (*i.e*., reducing type II error probability). Out of the initial 170 genes included in the analysis, 32 were differentially enriched (Table S1). Of these 32 genes, 7 were additionally assumed (via qualitative examination) to play a direct role in 3-CA degradation as identified in Table 1 and are discussed further.

**Table 1.**
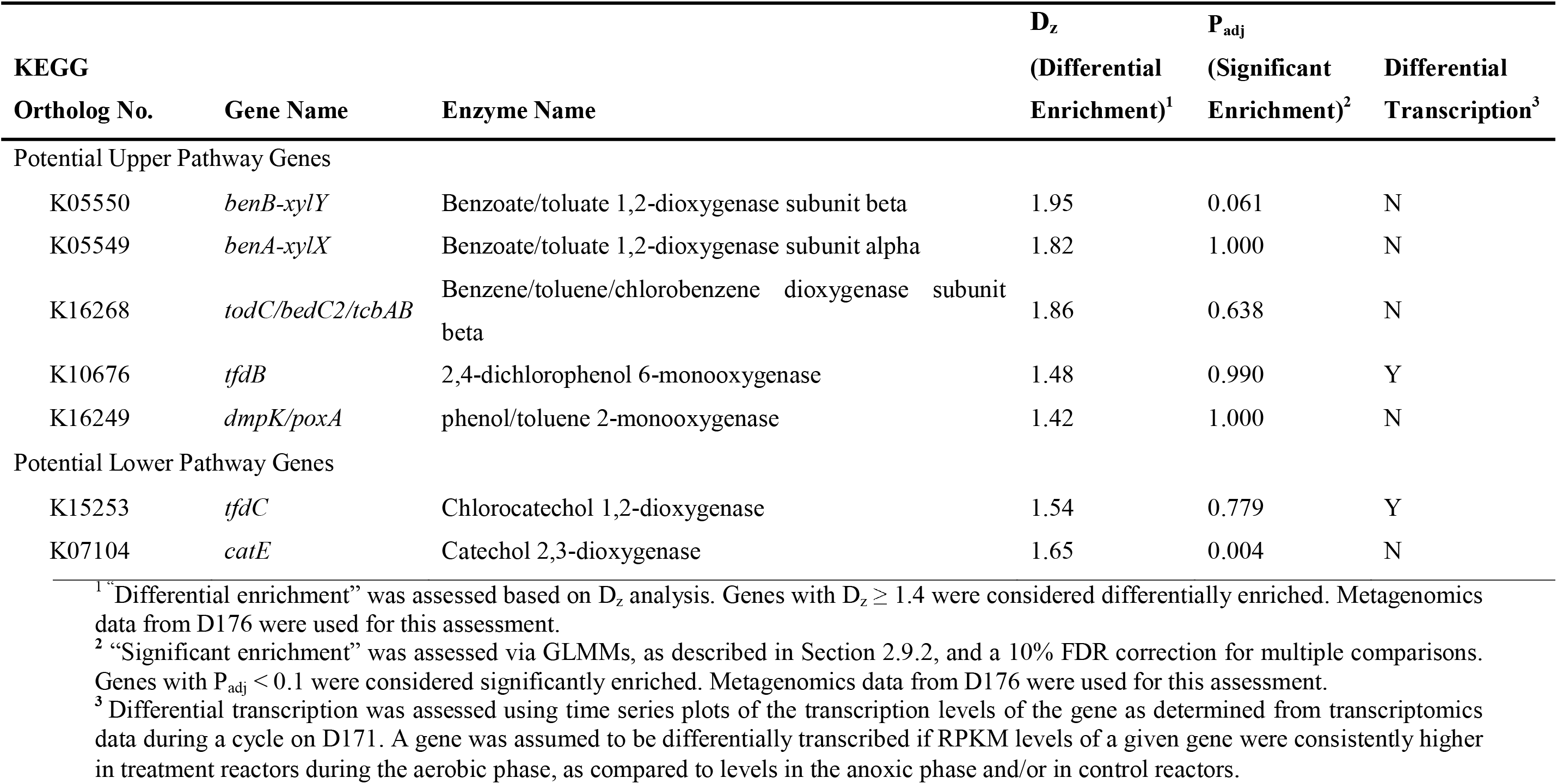
Major putative genes involved in 3-CA degradation as identified qualitatively from metagenomics and transcriptomics analyses. The full list of putative genes analysed is provided in Table S1.

To assess the significance of gene enrichment while also accounting for mean-variance relationship concerns (Wang et al., 2017), we performed a GLMM analysis on the full set of 170 genes to fit the data to a negative binomial distribution, using the *mvabund* package in R (v. 4.1.9). Both residuals vs fitted and mean-variance plots (available in Supplementary Files) supported the choice of a negative binomial distribution for the regression model. Univariate p-values for each of the 170 genes were adjusted for multiple comparisons using a step-down resampling procedure with a 10% P_adj_ cut-off, resulting in 11 genes being significantly enriched in treatment reactors. 9 of these 11 genes were also differentially enriched in treatment reactors (D_z_ ≥ 1.4) (Figure 4).

The significance of enrichment of each of the 170 genes is plotted against the levels of differential enrichment visually in Figure 4A, with the differentially enriched genes shown in greater detail in Figure 4B. Of the 7 differentially enriched genes that we qualitatively identified as being involved in 3-CA degradation (Table 1, Figure 4), 2 overlapped with the 9 identified above as being both significantly and differentially enriched (Table 1, Figure 4). All assembled MAGs were inspected to determine if any of these 7 genes were significantly enriched in treatment reactors, with negative results. The above differential enrichment (D_z_) process was repeated on MAG coverage between control and treatment reactors on D176, to determine the taxa relatively enriched in the treatment reactors, and potentially associated with 3-CA degradation (Table S2).

#### 2.9.3 Differential transcription of xenobiotics degradation genes from metatranscriptomics data

To determine whether the differentially enriched genes were also differentially transcribed in response to 3-CA input, we checked whether the transcription levels of a given gene related to 3-CA degradation rose during the aerobic phase of a cycle in treatment reactors compared to control reactors, and compared to the anoxic phase. The RPKM values for each relevant gene during a full cycle on D171 were plotted, including local polynomial regression fitting using the *loess* function from the *ggplot2* package (v.3.3.2) in R, including 95% confidence intervals.

## 3 Results and discussion

We investigated the effects of a sustained disturbance, caused by the presence of the xenobiotic compound 3-CA, on process performance, including COD removal, nitrification, and 3-CA persistence/degradation. Additionally, the advent of 3-CA degradation led to the identification of putative 3-CA degraders and their associated degradation pathways.

### 3.1 Impact of 3-CA on process performance

Nutrient profiles from all six reactors over several 12-h cycle studies revealed the impact of sustained 3-CA input on community function (Figures 1, 2 and S1 – S5). In the anoxic phase, control reactors consistently showed COD degradation (Figure 1B, S2B) and nitrate transformation (Figure 2C, S5B), with a transient nitrite spike (Figure 2B, S4B), indicating a complete denitrification pathway. Nitrification was also complete in control reactors in the aerobic phase, with ammonia levels falling to < 5 mg/L (Figure 2A, S3B) (apart from D110, where an inadvertent 3-CA input into reactor C3 caused a temporary loss of ammonia oxidation), an accompanying nitrite spike (Figure 2B, S4B), and a subsequent nitrate build-up (Figure 2C, S5B). In contrast, the treatment reactors initially experienced only partial COD degradation in the anoxic phase, which began to improve over time, as indicated by the change in COD profiles from D59 to D192, with COD levels on D124 not significantly different from those of the control reactors (Welch’s *t*-test P = 0.6) at the end of the cycle (Figure 1B, S2A). Ammonia oxidation was inhibited in treatment reactors in the aerobic phase (Figure 2A, S3A), with ammonia accrual over multiple consecutive cycles. However, some ammonia oxidation was observed in treatment reactors starting on D158 (Figure 2A, S3A), accompanied by some nitrite production in the aerobic phase (Figure 2B, S4A). These results indicated that COD degradation followed by ammonia oxidation gradually began to recover in treatment reactors, despite continuing 3-CA input.

Concurrent with COD degradation that developed from D95 onward in treatment reactors at the beginning of the aerobic phase (Figure 1B, S2A), 3-CA degradation was evident in all three treatment reactors (Figure 1A, S1), indicating that 3-CA accounted for the portion of COD degraded in treatment reactors during the aerobic phase. This strongly implies that treatment reactors had developed the ability to degrade 3-CA in response to sustained input. In turn, other processes like ammonia oxidation started to recover despite continued 3-CA input, as stated above. Batch experiments run separately (detailed in SI) showed that 3-CA was being degraded biologically (Figure S6).

The partial recovery of ammonia oxidation took about 35 days from the start of 3-CA input. Falk and Wuertz (2010) saw neither 3-CA degradation nor ammonia oxidation recovery during a 70-day experiment with sustained 3-CA input. However, they used a lower mixed liquor temperature (20° C versus 30° C in our experiment) and an inoculum from a wastewater treatment plant in a different region (temperate Davis, California, versus tropical Singapore) from a different sewershed (university campus, including multiple research facilities versus a mixed residential-industrial use in Singapore). Santillan et al. (2019) reported 3-CA degradation in a 35-day experiment with sustained 3-CA input, and partial ammonia oxidation recovery with less frequent 3-CA pulses. While their inoculum was from the same plant as ours, their reactors were much smaller, with a working volume of about 20 mL. Ma et al. (2020) reported full nitrification recovery within 12 days of a pulse 3-CA input (at lower concentrations) in a single-reactor study, possibly concurrent with the washout of 3-CA from the reactor. Kelly et al. (2004) reported a 17-day recovery period for total nitrification after a 1-chloro-2,4-dinitrobenzene pulse addition at a comparable concentration. The resumption of partial nitrification in our study may be attributed to the long SRT of 30 days. This allowed AOB to acclimate to the sustained 3-CA input. The onset of 3-CA degradation likely helped this recovery process. Both processes were aided by the source inoculum and the long SRT. A longer experimental run may have seen further recovery of ammonia oxidation and possibly of nitrite oxidation as well, but the aim of this experiment was to investigate the impact of 3-CA input on process performance and examine the mechanisms of established 3-CA degradation, for which the experimental timeline was deemed sufficient.

### 3.2 Organisms and pathways involved in 3-CA degradation

The emergence of 3-CA degradation in treatment reactors led to a search for the degrading organisms and their degradation pathway. Enrichment and isolation of 3-CA-degrading organisms using cultivation methods were unsuccessful (data not shown). We therefore turned to the molecular data collected – taxonomic data from amplicon sequencing, functional gene enrichment data from assembled metagenomics, and gene transcription data from metatranscriptomics – to evaluate potential 3-CA degraders and relevant functional genes.

#### 3.2.1 Organisms associated with 3-CA degradation

Three genera were significantly correlated with 3-CA degradation in all treatment reactors (after correction for multiple comparisons), namely *Gemmatimonas*, *OLB8* and *Taibaiella* (Figure 3). *Gemmatimonas* has been reported in a diverse range of activated sludge systems (Zhang et al., 2012; Ju et al. 2014). Santillan et al. (2019b) found this genus enriched under sustained 3-CA input conditions like those reported here. *OLB8* is not associated with aromatics degradation to our knowledge but belongs to the family *Saprospiraceae*, which is found in high-stress environments like low-temperature sludge (Johnston et al., 2019) and is associated with antibiotics resistance (Luan et al., 2020). Commonly found in activated sludge as a hydrolyser, it may break down complex organics (McIlroy and Nielsen, 2014; Szabó et al., 2017; Kondrotaite et al., 2022). The third genus, *Taibaiella*, has been found in phenol-degrading communities in microbial fuel cells (Shen et al., 2020), in petroleum-contaminated soil (Diallo et al., 2021), and in toxic landfill leachate (El-Fadel et al., 2018), indicating tolerance to xenobiotics, if not the ability to degrade them. Other ASVs that correlated (but not significantly) with 3-CA degradation included *Dokdonella*, *Burkholderia ottowia*, and two species of the order *Sphingobacteriales*. Sun et al. (2019) found *Sphingobacteriales* associated with aromatics degradation genes in phenol-degrading bioreactors. Other *Burkholderia* species degrade phenol (Ma et al., 2020b), and *Dokdonella* is associated with paracetamol (Palma et al., 2018) and polycyclic aromatic hydrocarbons degradation (Bacosa and Inoue, 2015).

Organisms associated with 3-CA degradation were also examined via differential enrichment of assembled MAGs in treatment reactors. In total, 60 medium quality (>80% completeness, <10% contamination) MAGs were recovered from all six reactors on D176 (Table S2). Of these, 11 were differentially enriched in treatment reactors (D_z_ ≥ 1.4) including the genera *JAAEKA01*, *Variibacter*, and *OLB8* and the species *Exiguobacterium_A indicum*. Of the differentially enriched MAGs, *OLB8* notably also correlated with 3-CA degradation in the amplicon sequencing data. This was the only organism identified as differentially enriched that correlated with 3-CA degradation in two independent datasets, making it a highly probable contender for 3-CA degradation. There were no other MAGs that matched the ASVs found to correlate with 3-CA degradation, likely because our MAG data represent only those taxa whose genomes could be assembled to an 80% completion threshold. It was expected that the amplicon sequencing and metagenomics data would not provide the same results, since they are two independent DNA-based methods of looking at microbial communities. The lack of complete overlap in results highlights the importance of applying multiple methods in molecular work.

Using molecular data to identify specific degraders as done in this study is not a common technique. Researchers attempting to identify degraders of a specific compound or members of a specific functional group are more likely to use stable isotope probing (Manefield et al., 2004; Aoyagi et al., 2018), or successive culturing techniques (for example, Chen et al., 2021). These methods may provide a more reliable means of identifying degrading organisms but involve the use of labelled isotopes in the feed to bioreactors, or successive rounds of culturing. The present study did not employ stable isotope probing because it was unclear at the onset of the experiment that 3-CA degradation would occur. Additionally, it is also unclear if 3-CA was used as a carbon and energy source, or as a nitrogen source. Cultivation methods were attempted but did not successfully enrich 3-CA degraders. The use of ‘omics data alone to identify degraders is less common, but allows for a faster, retroactive identification of potential degraders. However, it comes with its own drawbacks. It is possible that the taxa identified as potential 3-CA degraders in this study were merely able to adapt to 3-CA toxicity more efficiently than others. This would enable them to outcompete other organisms in terms of relative abundance. However, their correlation with the rate of 3-CA degradation makes it more likely that they were associated with the degradation of the compound.

#### 3.2.2 Differential enrichment of putative 3-CA degrading genes

3-CA is usually degraded by oxidation to a chlorocatechol (upper pathway) by oxidative deamination, followed by ring cleavage and subsequent degradation into tricarboxylic cycle intermediates (lower pathway) (Reineke, 1998; Bathe, 2004; Fuchs et al., 2011). We aimed to identify the upper pathway by searching for a mono- or dioxygenase, and the lower pathway via a ring cleavage enzyme. We acknowledge that 3-CA degradation may have occurred through a completely different and novel pathway. However, our investigation into degradation genes is confined to the most likely pathways for aromatics degradation identified in literature and known to occur in wastewater treatment systems (Hinteregger et al., 1992; Stockinger et al., 1992; Kaschabek et al., 1998).

Using assembled metagenomics data from D176, multiple upper and lower pathway genes were identified via the KEGG database (Aoki and Kanehisa, 2012), of which several were differentially enriched (and some also significantly enriched) in the treatment reactors (Figure 4A, Table S1). The enzyme involved in the first 3-CA oxidation step would theoretically be chloroaniline dioxygenase (Krol et al., 2012); unfortunately, its presence could not be validated because there is no KEGG ortholog for the gene encoding it. Also, chloroaniline dioxygenase may not have been present in the inoculum because it has a narrow substrate specificity (Krol et al., 2012) and may therefore not be common in activated sludge metagenomes. Additionally, the enzyme has previously been found encoded on a plasmid – this would prevent it from being identified via MAG analysis and may therefore be why it was not seen in our dataset. Other upper pathway genes were identified as putative contributors to 3-CA degradation, based on their identification in the 170-gene dataset and their ability to oxidize aromatic compounds (Table 1). These include benzoate/toluate 1,2-dioxygenase (KEGG ID K05550, *benB-xylY*; and K05549, *benA-xylX*), benzene/toluene dioxygenase (K16268, *tcbAB*), 2,4-dichlorophenol 6-monooxygenase (K10676, *tfdB*) and phenol/toluene 2-monooxygenase (K16249, *dmpK*). All of them were differentially enriched in treatment reactors (D_z_ ≥ 1.4) (Figure 4B), but only K05550 (*benB-xylY*) was significantly enriched (GLMM P_adj_ < 0.1) (Figure 4B, Table S1). They all normally act on their respective eponymous substrates. Whether chlorophenol enzymes can be induced by chloroaniline (via gene expression alone or enzyme production in the presence of the compound) is unclear. However, toluene dioxygenase degrades chloroanilines, including 3-CA (Nitisakulkan et al., 2014), and benzoate dioxygenase is associated with aniline degradation (Zhang et al., 2021). Furthermore, toluene dioxygenase has broad substrate specificity and can be induced by chloroanilines (Semak et al., 2012). It is also possible that the degradative enzymes were mediated by post-translational modifications, which implies that the DNA- and RNA-level data being used here may not pick them up. Thus, all these enzymes are potential candidates for initiating 3-CA degradation. Of the 9 genes that were both differentially and significantly enriched (Table S1), none other than benzoate/toluate 1,2-dioxygenase subunit beta (K05550, *benB-xylY*) were identified as an upper pathway dioxygenase. None of these genes were found within any of the MAGs assembled as part of this study.

For the potential lower degradation pathway, we focused on the relevant ring cleavage enzyme, because subsequent pathway segment genes are relatively common. The two major ring cleavage pathways for most chlorinated aromatics involve an ortho-cleavage or a meta-cleavage enzyme (Reineke, 1998). One enzyme of each type was differentially enriched in the treatment reactors (D_z_ ≥ 1.4) – a catechol 2,3-dioxygenase (K07104, *catE*; associated with meta-cleavage, potentially via 4-chlorocatechol) and a chlorocatechol 1,2-dioxygenase (K15253, *tfdC*; associated with ortho-cleavage, via 3- or 4-chlorocatechol) (Figure 4B, Table 1, Table S1). One of these enzymes (K07104, *catE*) was significantly and differentially enriched (GLMM P_adj_ < 0.1) (Figure 4B, Table S1). However, this enzyme is a catechol dioxygenase, not a chlorocatechol dioxygenase, and catechol dioxygenases are not usually induced by their chlorinated counterparts (Radianingtyas et al., 2003). Therefore, the metagenomics data did not conclusively identify which lower pathway gene may have contributed to 3-CA degradation. Of the 9 genes both differentially and significantly enriched (Figure 4, Table S1), most were identified as potential lower pathway genes, likely involved with post-cleavage pathway steps. The metagenomics data indicate that the most likely degradation pathway may have involved a meta-cleavage pathway via a benzoate/toluate dioxygenase (K05549, *benA-xylX*; K05550, *benB-xylY*) followed by a catechol dioxygenase (K07104, *catE*). We acknowledge that meta-cleavage pathways are less common for chlorinated aromatics than ortho-cleavage pathways (Reineke, 1998), but they have indeed been found (Kunze et al., 2009). It is also possible that the more relevant 3-CA degradation genes existed in the treatment reactors at levels below the detection limit of the metagenomics method. Additionally, the differential enrichment of one or more of the above pathway genes may be explained by gene hitchhiking (Oh et al., 2013), or by association with other degradative or 3-CA-tolerant genes that have not been identified here.

#### 3.2.3 Differential transcription of putative 3-CA degrading genes

Transcriptomics data from all six reactors on D171 were used to determine whether the upper and lower pathway genes identified above were transcribed in real time during a reactor cycle. 3-CA-degrading genes would likely be transcribed at higher levels in treatment reactors during the aerobic phase (when 3-CA removal occurred).

Four upper pathway genes were identified as likely contenders in Section 3.2.2. Of these, mean transcription levels of 2,4-dichlorophenol 6-monooxygenase (K10676; *tfdB*) were elevated in treatment reactors during the aerobic phase (Figure 5D). Of the two lower pathway genes identified earlier, chlorocatechol 1,2-dioxygenase (K15253; *tfdC*) also showed differential transcription in the treatment reactors during the aerobic phase (Figure 5B). Even though these genes were not significantly enriched (Table 1), their clear differential transcription makes them contenders for 3-CA degradation. Transcription levels of other major enzymes identified via metagenomics data as potential participants in 3-CA degradation (Table 1, Table S1) were not as clearly higher in treatment reactors during the aerobic phase. While potential intermediates like 3-chlorocatechol or 4-chlorocatechol were not measured in this experiment, the differential transcription of *tfdB* and *tfdC* in Figures 5D and 5B respectively indicates the formation of a complete pathway – the upper pathway, encoded by 2,4-dichlorophenol 6-monooxygenase (K10676, *tfdB*) showing differential transcription with the onset of aeration; and the lower pathway encoded by chlorocatechol 1,2-dioxygenase (K15253, *tfdC*) differentially transcribed toward the second half of the aerobic phase. Similarly, Sun et al. (2019) demonstrated differential transcription of phenol degrading genes in nitrifying activated sludge batch cultures via an ortho-cleavage pathway. Sato et al. (2019) also found differential enrichment and transcription of several aromatic degradation genes in response to heavy oil inputs to duplicate activated sludge reactors. However, all these studies used long-term sampling for gene transcription. Given the short half-life of mRNA (Selinger et al., 2003), mRNA profiles in a complex community like activated sludge can change rapidly. This study takes advantage of short-term sampling to consistently show the changes in mRNA levels of gene products related to a potential degradation pathway in the short term, within a short 12-hour period (one reactor cycle). Short-term mRNA studies for degradation dynamics are not common (for example, Leveau et al., 1999) and have not been done on an activated sludge microbial community, to our knowledge. Neither has a dichlorophenol monooxygenase been previously associated with chloroaniline degradation. This would be the first study to find an association between a chlorophenol monooxygenase and a chloroaniline degradation pathway.

**Figure 5.**
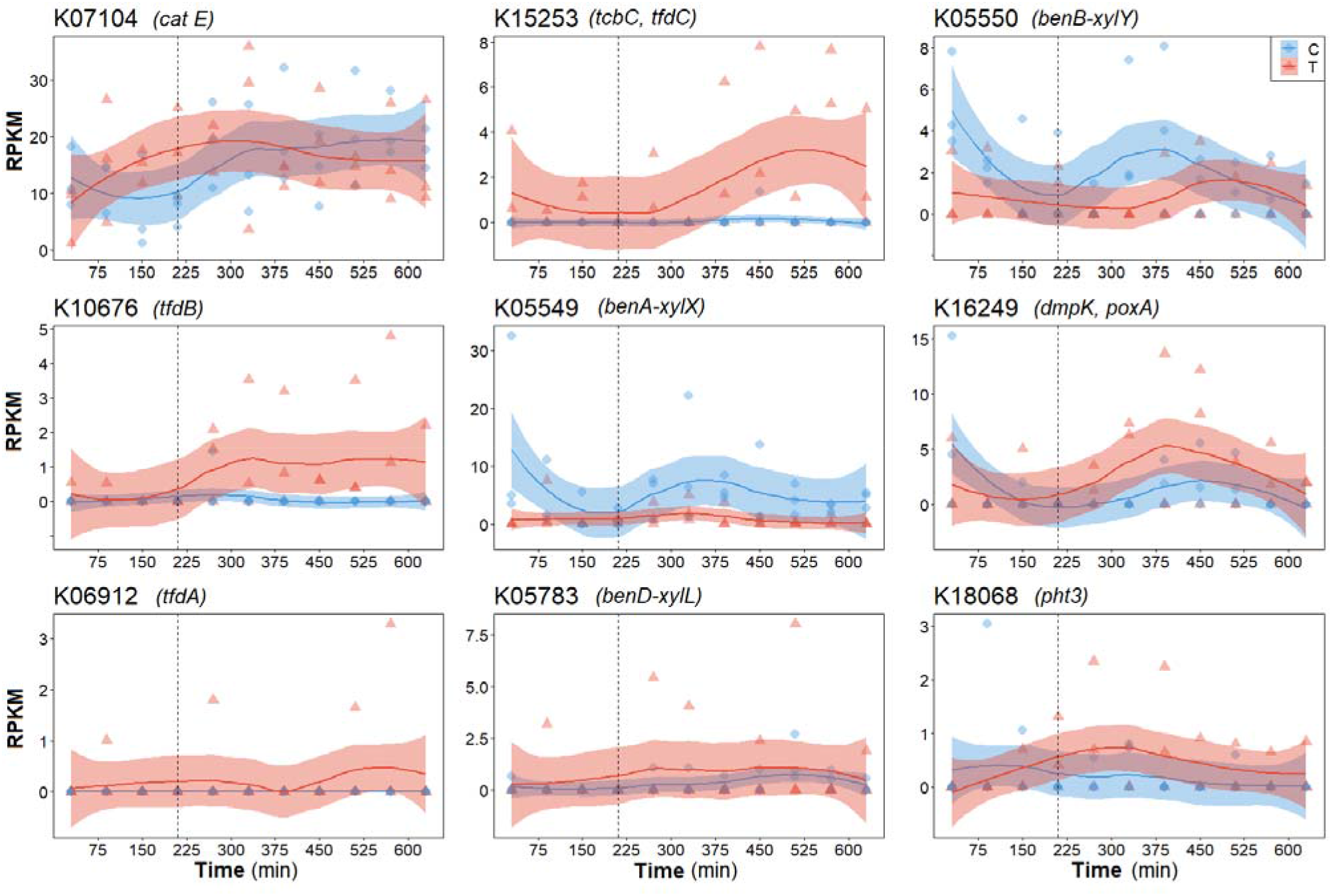
Average reads per kilobase million (RPKM) profiles of nine genes considered relevant to 3-CA degradation from which non-zero RPKM values were derived. RPKM profiles are shown over a single 12-h cycle on D171, averaged for the control (blue time series) and treatment (red time series) reactors for genes represented by KEGG orthologs K07104 (A), K15253 (B), K05550 (C), K10676 (D), K05549 (E), K16249 (F), K06912 (G), K05783 (H) and K18068 (I). The dotted vertical line (at x = 210 min) separates the anoxic and aerobic phases. Aeration began at 210 min. Lines display polynomial regression fitting, while shaded areas represent 95% confidence intervals.

#### 3.2.4 Full 3-CA degradation pathway

Combining the finding of differential transcription of both an upper and a lower pathway gene in the aerobic phase, together with process data showing 3-CA degradation in replicate reactors during the aerobic phase, two potential pathways for 3-CA degradation were identified. From the transcriptomics data, 3-CA degradation was found to occur via a modified ortho-cleavage pathway involving a dichlorophenol monooxygenase and a chlorocatechol dioxygenase. However, the metagenomics data present other genes that were differentially and significantly enriched in treatment reactors, suggesting a different, meta-cleavage pathway via a benzoate/toluate dioxygenase and a catechol dioxygenase.

It is likely that one or both pathways contributed to 3-CA degradation. While the metagenomics and transcriptomics data do not yield overlapping results, they add strength to the study, highlighting the importance of investigating multiple independent methods and using replicate reactors. However, it must be acknowledged that, although enriched in treatment reactors, the genes constituting the pathways identified above may not have contributed to 3-CA degradation, with an entirely different pathway or pathways responsible for actual degradation. It is worth noting that the use of known degrading genes from existing databases may help identify known pathways of degradation, but may not shed light on novel or less known pathways.

Since the impacts of 3-CA and its subsequent degradation were clearly seen for the nitrification process, it is likely that 3-CA had a major impact on AOB and NOB communities. The changes in nitrifier communities and nitrification genes merit further study.

## 4 Conclusions

In this study, we ran a controlled experiment to investigate the impact of sustained 3-chloroaniline input on replicated wastewater-treating bioreactors from a process as well as microbial community perspective.

- Full removal of 3-CA was observed in all three treatment reactors (after a three-week period of sustained input) with the recovery of ammonia oxidation, which was initially inhibited by 3-CA input.
- Using a combination of metagenomics and metatranscriptomics, two putative pathways for 3-CA degradation were identified:

- The first putative pathway involved a phenol monooxygenase and a subsequent ortho-cleavage enzyme. These genes were both differentially transcribed during the aerobic phase of a cycle and differentially enriched (albeit not significantly).
- The second putative pathway included a benzoate/toluate dioxygenase followed by a meta-cleavage enzyme. These genes were both differentially and significantly enriched in the treatment reactors.
- Amplicon sequencing and assembled metagenomics data showed multiple taxa that correlated significantly with 3-CA degradation and were differentially enriched in treatment reactors, respectively. The genus *OLB8* was the only taxon identified in both datasets in this regard, making it a possible actor in 3-CA degradation.

## Acknowledgements

This work was funded by the Singapore Ministry of Education and the National Research Foundation under an RCE grant awarded to the Singapore Centre for Environmental Life Sciences Engineering (SCELSE). The authors would like to thank Angel Anisa Cokro and Angel Anika Cokro, Anita Ng and Sean Chew for their help with reactor operation and sample processing; Larry Liew Chee Wai for his help with sample collection and storage; Daniela Drautz-Moses and colleagues from the SCELSE sequencing core facility for obtaining short read sequencing data used in this study; Krithika Arumugam for data processing and bioinformatics; and Siao Yun Chang and colleagues at Public Utilities Board for providing access to sludge as inoculum.

## Author Contributions

SW and HS conceived the study and SW obtained the funding; HS designed the study and performed the experiments; and HS and ES performed the molecular work to prepare samples for sequencing. USCS developed the RNA extraction protocol and extracted RNA. ES did the bioinformatic analysis on the 16S rRNA gene amplicon sequences; FC performed metagenomics assembly bioinformatics; and RW analysed the transcriptomics data. HS and ES interpreted individual data sets; HS wrote the manuscript; and the other authors contributed to individual sections and edited the draft.

## Data Availability

DNA sequencing data are available at NCBI BioProjects under accession numbers PRJNA720804 and PRJNA720805 (upload in progress). See Supplementary Information (SI) for details about the 3-CA degradation batch experiment, the calculation of 3-CA degradation rates in treatment reactors, a list of genes associated with xenobiotics degradation that were differentially enriched in treatment reactors, a list of medium-quality MAGs, and chemical profiles over single 12-h cycles for all reactors measured on ten different days of the study. All other relevant data and statistics related to metagenomics, MAGs, metatranscriptomics, ASVs, and process performance are publicly accessible on FigShare (https://doi.org/10.6084/m9.figshare.20130503).

## Competing Interests

The authors declare no competing interests.

## Notes

### Competing Interest Statement

The authors have declared no competing interest.

### Summary of Updates

Improved manuscript introduction and discussion. Added Generalized linear mixed models (GLMMs) and metagenome-assembled genomes (MAGs) to the analysis.

