## Supplementary Information for "Metagenomics and Metatranscriptomics Suggest Pathways of 3-Chloroaniline Degradation in Wastewater Reactors"

### S1 Methods

#### *S1.1 Additional Details on Experimental Design*

The synthetic wastewater fed to all six reactors in the feed phase of each cycle during the acclimation period was adapted from Hesselmann et al. (1999) and contained (mg/L in mixed liquor in each reactor after feeding and being diluted in mixed liquor): sodium acetate (112.5), dextrose (45), yeast extract (67.5), soy peptone (60), meat peptone (60), casein peptone (90), urea (15), ammonium bicarbonate (90), ammonium chloride (169), disodium hydrogen phosphate (720), potassium dihydrogen phosphate (130), calcium chloride dihydrate (10.5) and magnesium sulphate heptahydrate (112.5). The medium also contained 2 mL of the unaltered trace element stock (Hesselmann et al.,1999) per litre of medium. During acclimation and in control reactors, the first six macrocomponents contributed to the COD, amounting to about 500 mg/L in the mixed liquor; the next three (ammonium-based) components contributed to the total nitrogen concentration of about 70 mg N/ L; and the phosphates were used to buffer the medium and maintain a pH around 7.5 to facilitate nitrification. The medium

Water quality parameters were measured as described by Santillan et al. (2021). Standard nutrient analysis techniques per Standard Methods (APHA-AWWA-WEF, 1998) were used, targeting COD (Standard Methods 5220 D) and nitrogen species (ammonium, nitrite and nitrate) using ion chromatography (Standard Methods 4500-NH3 for ammonium; 4110 B for nitrate and nitrite). 3-CA was measured on a Shimadzu Prominence high pressure liquid chromatography (HPLC) system (Shimadzu Corporation, Kyoto, Japan) equipped with a UV-VIS PDA detector using an Ascentis C18 5-μm column (Sigma-Aldrich, St. Louis, Missouri, USA). An isocratic 50:50 water:methanol solvent was used at a flow of 0.3 mL/min, and 3-CA peaks were identified and measured at 199, 237 and 286 nm.

Genomic DNA was extracted from samples collected from bioreactors (500 μL of sludge per sample) using the FastDNA Spin Kit for Soil (MP Biomedicals) with alterations to the manufacturer’s protocol to increase DNA yield as detailed elsewhere (Santillan et al., 2019).

Extracted DNA (using the FastDNA Spin Kit for Soil (MP Biomedicals-) from D0, D56, D63, D112 and D164 was amplified for the 16S rRNA gene in triplicate 50-μL PCR reactions using primer set 530f/U1053r targeting the V3~V4-V5 variable regions of the bacterial 16S rRNA gene (Thijs et al., 2017). Each 50-μL PCR reaction contained 25 μL ImmoMix reagent (Meridian Bioscience, Cincinnati, OH, USA) and 200 ng of DNA extract and nuclease-free water. The PCR program included initial denaturation at 95º C for 10 min, followed by 30 cycles of denaturation (95ºC, 1 min), annealing (58ºC, 30 s) and extension (72ºC, 1 min); and a final extension at 72º C for 7 min. The triplicate PCR amplicons from each sample set were then pooled and purified using the QIAquick PCR purification kit (Qiagen), with a slightly altered protocol as described in Santillan et al. (2021). The purified amplicons were then inspected for quality using agarose gels and quantified using a Qubit 3.0 fluorometer (ThermoFisher Scientific, Waltham, MA, USA).

The libraries were sequenced in-house at the Singapore Centre on Environmental Life Sciences Engineering (SCELSE) on an Illumina MiSeq (v.3) platform with 20% PhiX spike-in, at 300 bp paired-end read-length. Sequenced sample libraries were processed with the *dada2* (v.1.3.3) R-package (Callahan et al., 2016), allowing inference of amplicon sequence variants (ASVs) (Callahan et al., 2017). Illumina adaptors and PCR primers were trimmed prior to quality filtering. Sequences were truncated after 280 and 255 nucleotides for forward and reverse reads, respectively. After truncation, reads with expected error rates higher than 3 and 5 for forward and reverse reads, respectively, were removed. Since paired-end reads could not be merged with a minimum overlap of 20 bp, only forward reads were kept for further analysis. Chimeric sequences (0.3% on average) were identified and removed. For a total of 30 samples, an average of 48,659 reads were kept per sample after processing, representing 61% of the average forward input reads. Taxonomy was assigned using the SILVA database (v.132) (Glöckner et al., 2017). Samples were normalized to the sample with the lowest total read count after processing (18,353). To determine the impacts of 3-CA input on the nitrifier fraction of the microbial community, ASVs identified as representing ammonia oxidizing bacteria and nitrite oxidizing bacteria were extracted from the matrix containing all ASVs and their relative abundances were calculated as percentages of total normalized counts per sample. Whole microbial community-level analysis of this data is available elsewhere (Santillan et al., 2021).

*S1.1 Batch experiment on 3-CA degradation*

To determine whether 3-CA was being metabolized biologically, a batch experiment was performed using sludge samples from each reactor on D170. The batches were run in 500 mL baffled flasks, in triplicate. In each of these flasks, 100 mL of sludge was added from a single reactor and topped up with 100 mL of synthetic wastewater containing 3-CA. This amounted to a total batch volume of 200 mL. Three such flasks were run with sludge from each reactor, amounting to a total of 18 batch cultures. These batch cultures were accompanied by three abiotic control flasks containing 200 mL of deionised water spiked with 3-CA to the same concentration of 70 mg/L. The flasks were placed in shaking incubators at 30$^{o}$ C and 150 rpm to stay mixed and aerated over a 24-h period. At the beginning and end of this period, about 1 mL of mixed liquor was taken from each flask, filtered through 0.2-μm filters and analyzed by HPLC for 3-CA concentrations as described below.

*S1.2 Batch experiment data analysis*

3-CA concentrations measured using HPLC from the sludge samples taken from each flask described in Section 2.4 from before and after the batch experiment were used to calculate the percent of total input 3-CA remaining after 24 hours. The mean and standard deviation of the mean 3-CA level remaining in three replicate batch cultures from each source reactor were calculated. A *t*-test was used to determine the differences among the batches from each of the six source reactors.

*S1.3 3-CA degradation rates*

To determine degradation rates of 3-CA in the treatment reactors, data from the cycle study on D171 were used. 3-CA degradation rates from other cycle studies where degradation had stabilised in all three reactors were not found to be different. 3-CA concentrations measured during the aerobic phase, where 3-CA degradation primarily occurred, were used to calculate the first-order decay coefficient, using the time elapsed and the natural logarithm of the measured 3-CA concentration. Data were included throughout the aerobic phase, to account for the initial period of fast degradation, and the subsequent period of slow degradation. Only cycle studies past D95, after stable 3-CA degradation was seen in all treatment reactors, were used to calculate degradation rates.

#### *S1.4 Inadvertent 3-CA Addition to Control Reactor*

3-CA was unintentionally added to control reactor C3 during one cycle. This occurred at the beginning (feed phase) of a single 12-h cycle on D99. This unintentional input was at the same 3-CA concentration as that fed at every cycle to the treatment reactors.

### S2 Results and Discussion

#### *S2.1 3-CA degradation rates*

Specific 3-CA degradation rates calculated toward the end of the experiment, once 3-CA degradation had stabilized (D158 and beyond), averaged 7.5 mg 3-CA $g^{-1}$ VSS $h^{-1}$ which is similar to the rates seen by Zhu et al. (2012) for 3-CA degradation in batch cultures from aerobic granular sludge at similar input concentrations. The mean first-order kinetics constant on D171 was about 0.02 $min^{-1}$, five orders of magnitude higher than those reported by Susarla et al. (1997) for multi-chlorinated anilines in anaerobic sediment. This is likely because aerated bioreactors operated at 30ºC are likely to bring about faster degradation than microbial activity in natural sediments. Bathe et al. (2009) reported much lower non-specific degradation rates, on the order of 1.5 mg $L^{-1}$ $h^{-1}$ in suspended biofilm cultures; however, their cultures had been previously acclimated to varying loads of 3-CA input and were subjected to much higher 3-CA concentrations of about 200 mg/L in mixed liquor.

#### *S2.2 Batch 3-CA degradation tests*

Studies to test the 3-CA degradation ability of the six bioreactor sludges were conducted in separate batch cultures using sludge from each of the six reactors, on D170. This experiment lasted 12 h, or one bioreactor cycle. This batch study showed almost complete 3-CA removal from all three treatment-reactor sludges, while negligible 3-CA degradation occured (comparable to the abiotic control) in the three control reactors (Figure S6). There were significant differences in the amount of 3-CA remaining in batch cultures of sludges from control and treatment reactors (*t*-test $P = 2.04 \times 10^{-7}$). Sludges from reactors T1, T2 and T3 removed similar amounts of 3-CA, with nearly 0% 3-CA remaining, while sludges from reactors C1, C2 and C3 were similar, with greater than 90% of 3-CA remaining in the cultures (Figure S6). This batch experiment, conducted separately from the bioreactor study, demonstrated that 3-CA was being processed biologically in the treatment reactor sludges.

#### *S2.3 Inadvertent 3-CA input to Reactor C3*

Control reactor C3 was subjected to an unintentional pulse loading of 3-CA on D99, at the same concentration as the treatment reactors. While this was not part of the experimental plan, it provided insight into the response of the microbial community to a shock 3-CA load. COD removal, denitrification and ammonium oxidation in C3 were disrupted for less than three weeks after the pulse input; however nitrite oxidation suffered the most, and nitrate production took almost five weeks to recover to pre-pulse levels. These results are incorporated into the calculation of average nutrient profiles in control reactors Figures S2B, S3B and S4B, on D110, D124 and D136. Particularly evident is the higher average ammonia profile on D110 (Figure S3B), which is the direct result of the loss of ammonia oxidation in reactor C3 due to this pulse input on D99. Earlier studies that examined shock 3-CA loadings (Boon et al., 2003; Marzorati et al., 2013) found no nitrification recovery following a shock load, but these studies were run for only about two weeks, much less than the normal doubling time of nitrifiers (Painter, 1970). More recently Ma et al. (2020) saw nitrification recover in about two weeks after a pulse 3-CA input. This is more in keeping with our results. Nitrite oxidation took longer to recover in our reactors possibly because of the reasons covered in Section 3.2.

The results presented in the main body of this manuscript primarily show results from days before this inadvertent input to reactor C3 occurred, and after its impacts on process had recovered.

### Supplemental Figures and Figure Legends

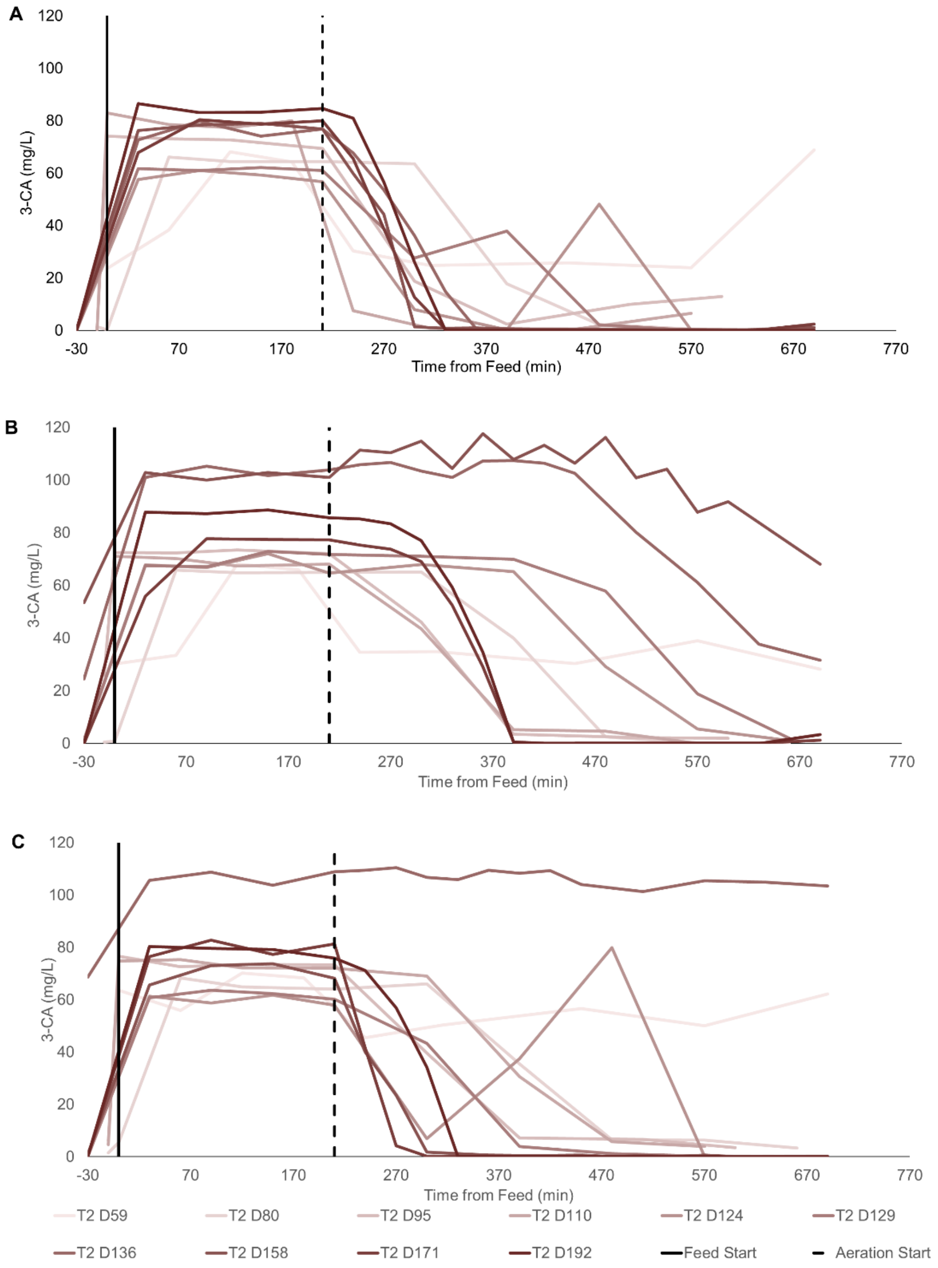


**Figure S1.** 3-CA concentration profiles in each of the treatment reactors on ten distinct days of the experiment showing degradation of COD and 3-CA over single 12-h cycles for all six reactors. The x-axes represent time from the commencement of feeding in min. The dotted vertical line (at x = 210 min) separates the anoxic and aerobic phases. Aeration began at 210 min. The y-axes represent 3-CA, mg/L in **(A)** Reactor T1, **(B)** Reactor T2 and **(C)** Reactor T3.

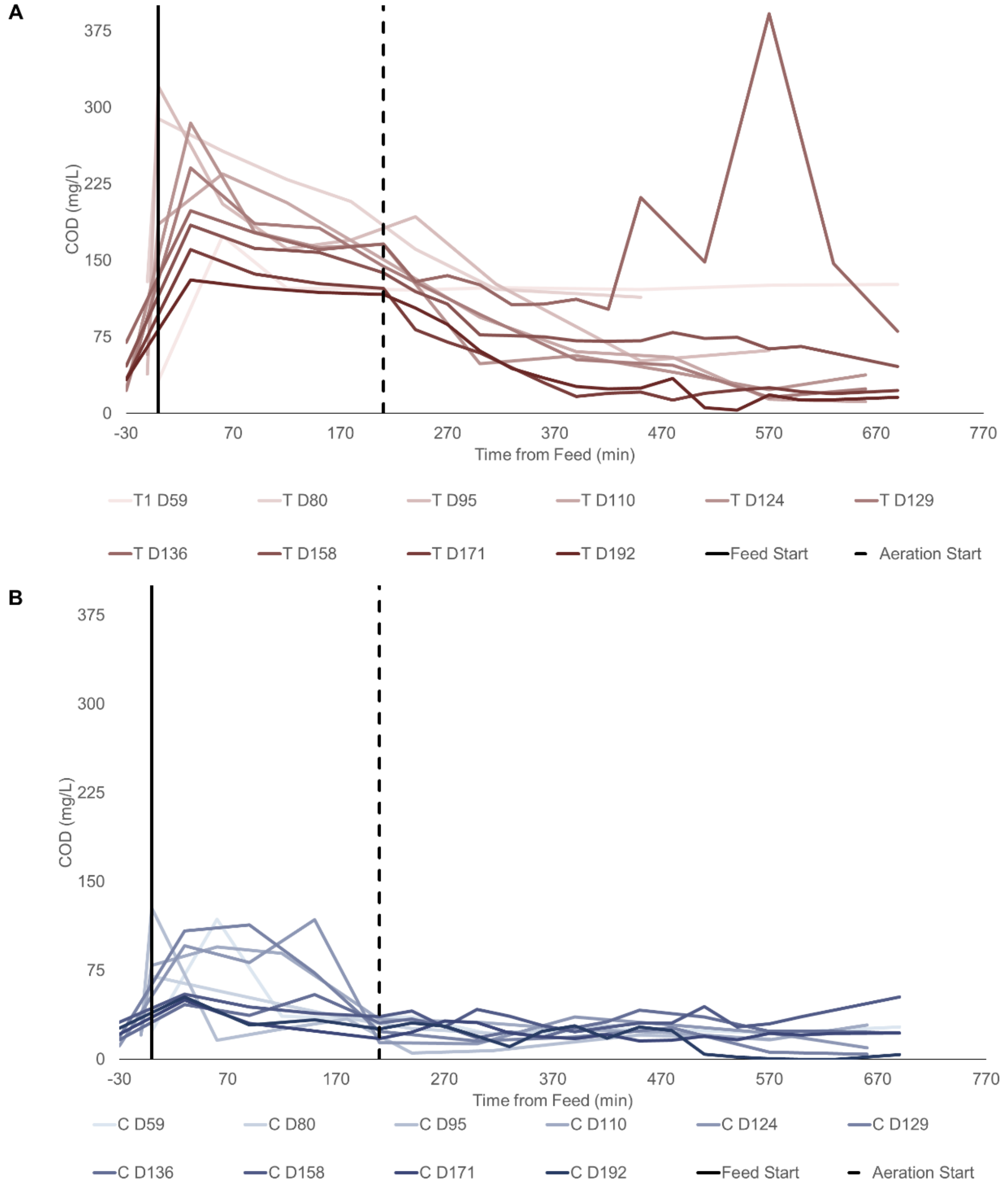


**Figure S2**. Average COD concentration profiles from ten distinct days of the experiment showing degradation of COD over single 12-h cycles for all six reactors. The x-axes represent time from the commencement of feeding in min. The dotted vertical line (at x = 210 min) separates the anoxic and aerobic phases. Aeration began at 210 min. The y-axes represent the mean COD, mg/L, in (**A**) Treatment Reactors (T1, T2 and T3) and (**B**) Control Reactors (C1, C2 and C3).

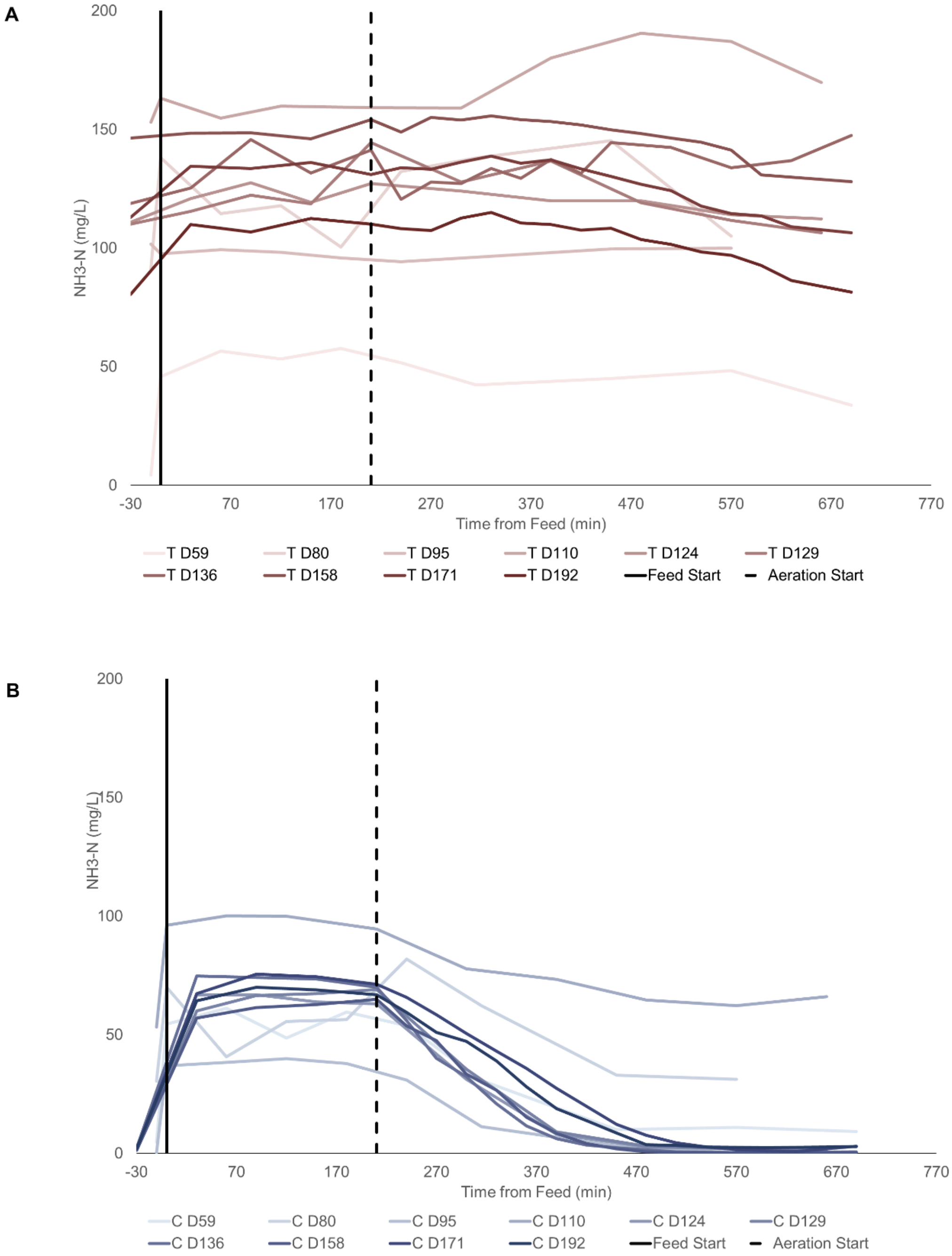


**Figure S3**. Average ammonia-nitrogen concentration profiles from ten distinct days of the experiment showing degradation of $NH_3$-N over single 12-h cycles for all six reactors. The x-axes represent time from the commencement of feeding in min. The dotted vertical line (at x = 210 min) separates the anoxic and aerobic phases. Aeration began at 210 min. The y-axes represent the mean $NH_3$-N, mg/L, in (**A**) Treatment Reactors (T1, T2 and T3) and (**B**) Control Reactors (C1, C2 and C3).

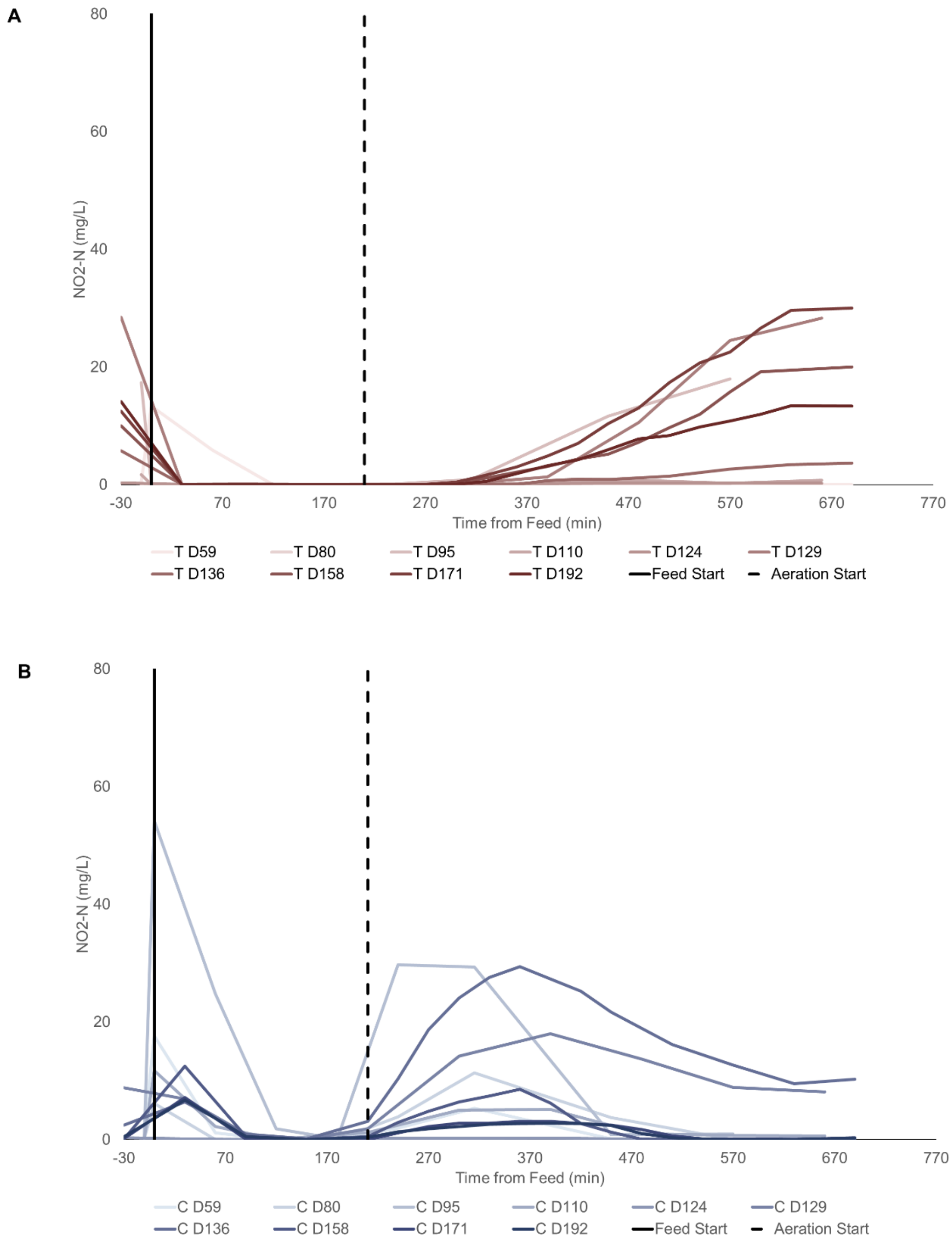


**Figure S4**. Average nitrite-nitrogen concentration profiles from ten distinct days of the experiment showing degradation of $NO_2$-N over single 12-h cycles for all six reactors. The x-axes represent time from the commencement of feeding in min. The dotted vertical line (at x = 210 min) separates the anoxic and aerobic phases. Aeration began at 210 min. The y-axes represent the mean $NO_2$-N, mg/L, in (**A**) Treatment Reactors (T1, T2 and T3) and (**B**) Control Reactors (C1, C2 and C3).

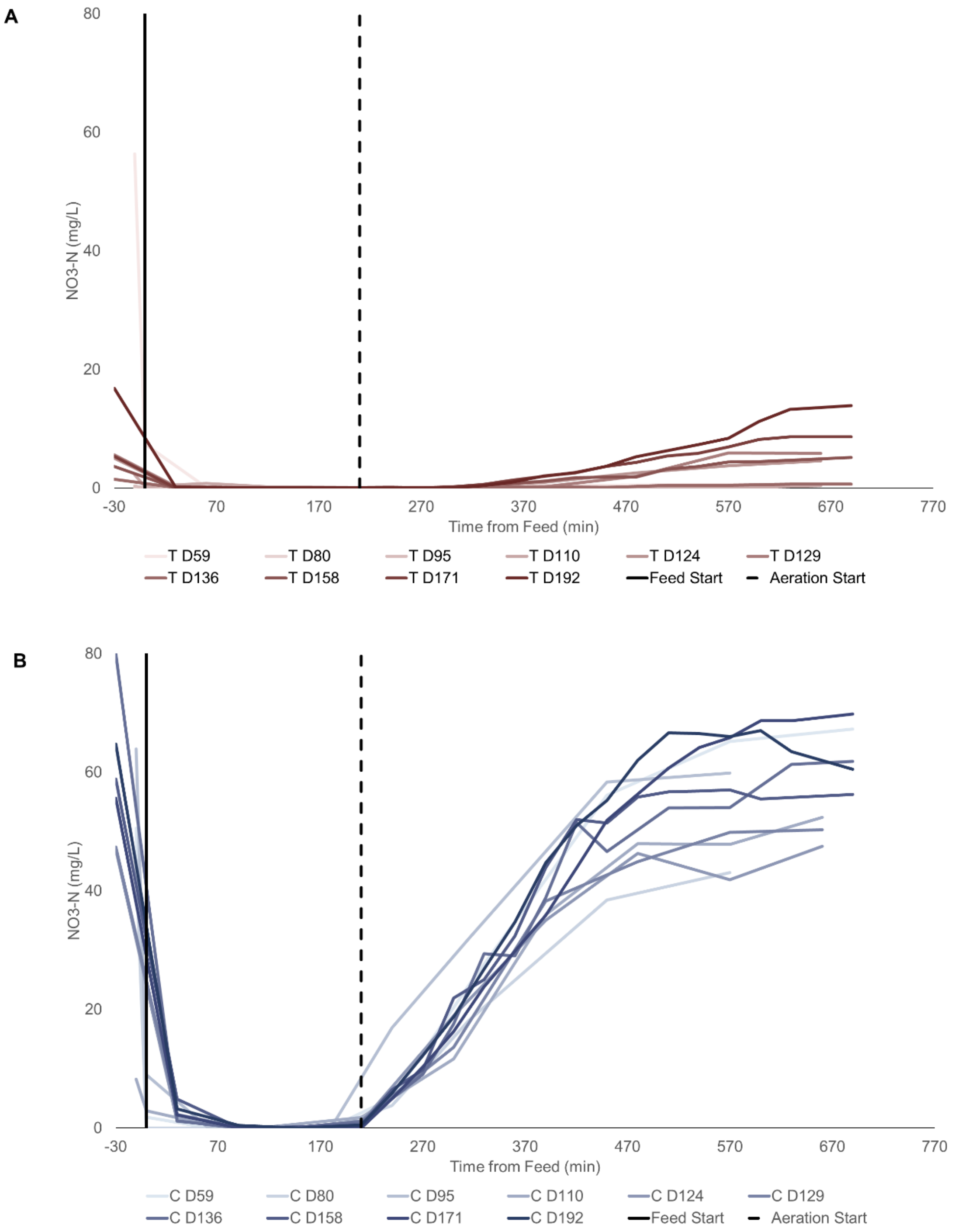


**Figure S5**. Average nitrate-nitrogen concentration profiles from ten distinct days of the experiment showing degradation of $NO_3$-N over single 12-h cycles for all six reactors. The x-axes represent time from the commencement of feeding in min. The dotted vertical line (at x = 210 min) separates the anoxic and aerobic phases. Aeration began at 210 min. The y-axes represent the mean $NO_3$-N, mg/L, in (**A**) Treatment Reactors (T1, T2 and T3) and (**B**) Control Reactors (C1, C2 and C3).

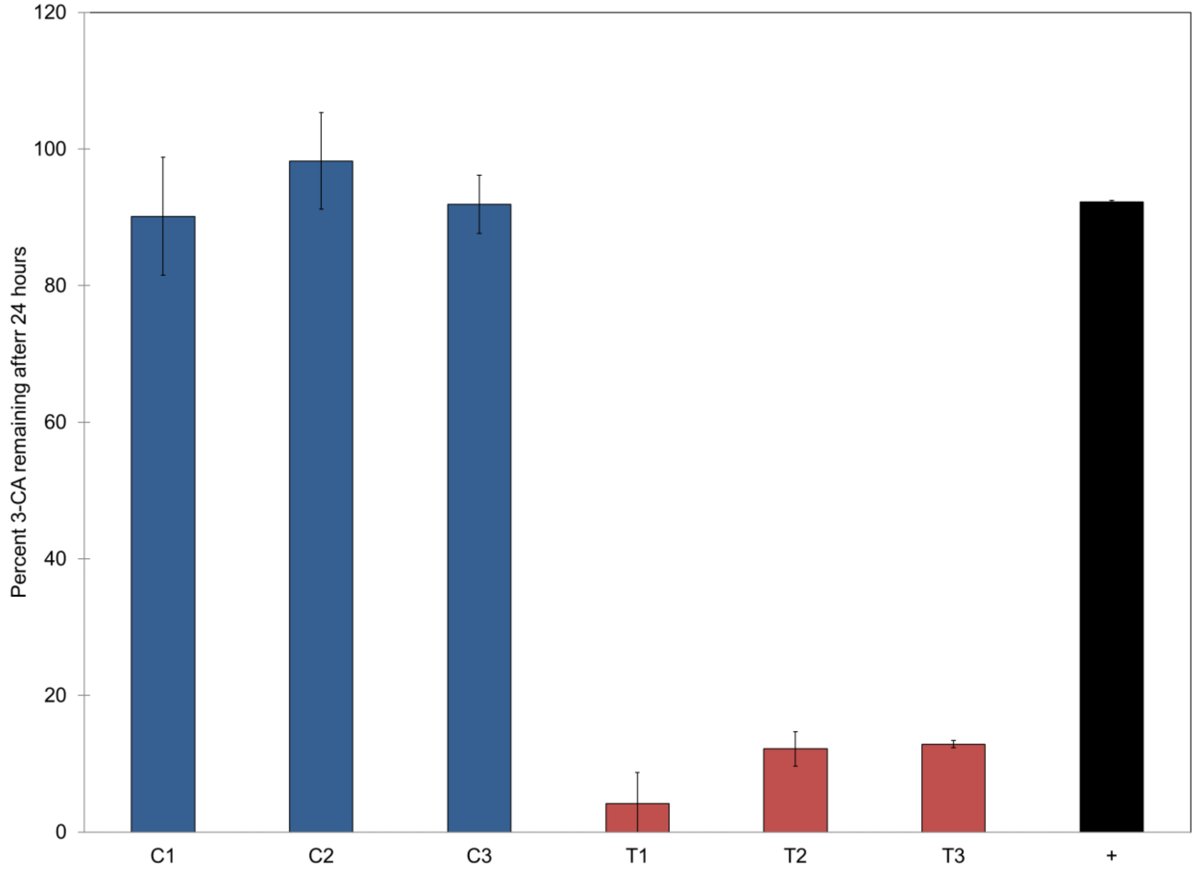


**Figure S6.** Percent of total input 3-CA remaining after 12 h of shaking in triplicate batch cultures of sludge taken from each bioreactor on D170. Bars show the mean 3-CA remaining over three replicate batch cultures from each source reactor, while error bars represent one standard deviation of the mean. Blue bars represent sludge from control reactors and red bars sludge from treatment reactors. The black bar refers to the positive control with no sludge.

### Supplemental Tables and Table Legends

**Table S1**. Differentially enriched genes ($D_z \geq 1.4$) associated with xenobiotics degradation that were identified in metagenomics data from D176 in all six reactors, along with their respective significance of enrichment. Putative 3-CA degradation pathway genes and differentially transcribed genes are also identified.

| S No. | KEGG ID | Differential Enrichment ($D_z \geq 1.4$) | Significant Enrichment (GLMM) $P_{adj}$ [1] | Potential Pathway Gene?[2] | Differential Transcription?[3] |
|---|---|---|---|---|---|
| 1 | K05550 | 1.95 | **0.060** | U | N/A |
| 2 | K16268 | 1.86 | 0.635 | U | N/A |
| 3 | K15237 | 1.85 | 0.281 | N | N/A |
| 4 | K18069 | 1.85 | **0.080** | N | N/A |
| 5 | K14578 | 1.84 | 0.323 | N | N/A |
| 6 | K05549 | 1.82 | 1.000 | L | No |
| 7 | K07823 | 1.78 | 0.879 | N | N/A |
| 8 | K04105 | 1.77 | 1.000 | U | No |
| 9 | K11947 | 1.77 | 1.000 | U | No |
| 10 | K01512 | 1.73 | **0.004** | N | N/A |
| 11 | K06912 | 1.72 | 0.999 | N | No |
| 12 | K00626 | 1.69 | **0.004** | U | No |
| 13 | K01782 | 1.69 | **0.004** | N | N/A |
| 14 | K04117 | 1.67 | 0.327 | N | N/A |
| 15 | K10216 | 1.67 | **0.004** | U | No |
| 16 | K01101 | 1.66 | **0.004** | L | Yes |
| 17 | K07104 | 1.65 | **0.004** | U | N/A |
| 18 | K07535 | 1.63 | 1.000 | N | N/A |
| 19 | K07536 | 1.62 | 1.000 | U | Yes |
| 20 | K03863 | 1.60 | 1.000 | N | N/A |
| 21 | K04116 | 1.60 | 1.000 | U | Yes |

| | | | | | |
|---|---|---|---|---|---|
| 22 | K10622 | 1.60 | 0.148 | N | N/A |
| 23 | K18249 | 1.59 | 0.843 | U | No |
| 24 | K15253 | 1.54 | 0.772 | U | No |
| 25 | K05783 | 1.54 | 0.996 | N | N/A |
| 26 | K10676 | 1.48 | 0.988 | N | N/A |
| 27 | K01501 | 1.48 | 1.000 | N | N/A |
| 28 | K00632 | 1.47 | 0.400 | N | N/A |
| 29 | K18068 | 1.44 | 1.000 | N | N/A |
| 30 | K01692 | 1.43 | **0.040** | N | N/A |
| 31 | K16249 | 1.42 | 1.000 | U | No |
| 32 | K01612 | 1.42 | 1.000 | N | N/A |

**Notes:**

1. Adjusted P-values using a step-down resampling procedure, from univariate GLMM analysis of the list of 170 genes recovered from the KEGG metabolic pathways within the xenobiotics biodegradation and metabolism category. In bold, adjusted P-values below 10% cutoff. None of these genes were found to be more abundant in assembled MAGs that were differentially enriched in treatment reactors (Table S2). Full gene list and adjusted univariate P-values are available as supplementary information.
2. U = potential upper pathway gene; L = potential lower pathway gene; N = not considered a major pathway gene
3. Differential transcription assessed by higher levels of transcription seen from transcriptomics data in treatment reactors in the aerobic phase. N/A implies transcription data was not processed for the given gene.

**Table S2**. List of medium-quality metagenome-assembled genomes across all reactors in D176.

| **Taxonomy from GTDB (Genome Taxonomy Database)** [*Bacteria* domain] | **Enrich**[1] ($D_z$) | **Length** (bp) | **GC** (%) | **Contigs** | **Comp**[2] (%) | **Cont**[3] (%) |
|---|---|---|---|---|---|---|
| p__Chloroflexota;c__Anaerolineae;o__Caldilineales;f__JAAEKA01;g__JAAEKA01;s__ | 2.00 | 3669767 | 65.99 | 711 | 82.33 | 4.36 |
| p__Proteobacteria;c__Alphaproteobacteria;o__Rhizobiales;f__Xanthobacteraceae;g__Variibacter;s__ | 1.89 | 3086568 | 65.65 | 128 | 82.68 | 1.01 |
| p__Firmicutes;c__Bacilli;o__Exiguobacterales;f__Exiguobacteraceae;g__Exiguobacterium_A;s__Exiguobacterium_A indicum | 1.79 | 3252289 | 47.6 | 87 | 98.28 | 3.45 |
| p__Planctomycetota;c__Planctomycetes;o__Isosphaerales;f__Isosphaeraceae;g__;s__ | 1.68 | 5698706 | 66.66 | 748 | 80.86 | 3.65 |
| p__Chloroflexota;c__Anaerolineae;o__SBR1031;f__A4b;g__OLB15;s__OLB15 sp001567085 | 1.67 | 3991379 | 54.56 | 14 | 97.52 | 3.36 |
| p__Bacteroidota;c__Bacteroidia;o__Chitinophagales;f__Saprospiraceae;g__OLB8;s__OLB8 sp001567405 | 1.57 | 3777414 | 41.25 | 120 | 98.46 | 0.51 |
| p__Bacteroidota;c__Bacteroidia;o__Chitinophagales;f__Chitinophagaceae;g__Flavipsychrobacter;s__ | 1.51 | 2969905 | 41.46 | 251 | 99.23 | 1.59 |
| p__Bacteroidota;c__Kapabacteria;o__Kapabacteriales;f__Kapabacteriaceae;g__Kapabacteria;s__Kapabacteria thiocyanatum | 1.51 | 3321029 | 59.63 | 32 | 98.28 | 0.00 |
| p__Bacteroidota;c__Bacteroidia;o__NS11-12g;f__UKL13-3;g__;s__ | 1.46 | 2320298 | 33.53 | 42 | 94.31 | 1.03 |
| p__Bacteroidota;c__Bacteroidia;o__Chitinophagales;f__Chitinophagaceae;g__Palsa-955;s__ | 1.43 | 3681358 | 42.56 | 79 | 99.49 | 1.54 |
| p__Armatimonadota;c__UBA5829;o__DSUL01;f__DSUL01;g__;s__ | 1.41 | 4141864 | 57.78 | 186 | 100 | 3.45 |
| p__Bacteroidota;c__Bacteroidia;o__Chitinophagales;f__Chitinophagaceae;g__;s__ | 1.39 | 2437211 | 44.42 | 20 | 98.21 | 0.00 |
| p__Bacteroidota;c__Bacteroidia;o__AKYH767-A;f__OLB10;g__OLB10;s__OLB10 sp001567275 | 1.34 | 3090274 | 39.52 | 235 | 98.46 | 1.43 |
| p__Chloroflexota;c__Chloroflexia;o__Thermomicrobiales;f__UBA6265;g__UBA6265;s__ | 1.33 | 3357120 | 57.93 | 812 | 91.38 | 6.97 |
| p__Proteobacteria;c__Gammaproteobacteria;o__Burkholderiales;f__Burkholderiaceae;g__SCN-69-89;s__SCN-69-89 sp003577305 | 1.28 | 3305121 | 69.45 | 374 | 91.95 | 7.76 |
| p__Proteobacteria;c__Alphaproteobacteria;o__Rhizobiales;f__Xanthobacteraceae;g__62-47;s__ | 1.27 | 4102035 | 65.45 | 854 | 84.23 | 4.81 |
| p__Gemmatimonadota;c__Gemmatimonadetes;o__Gemmatimonadales;f__Gemmatimonadaceae;g__SCN-70-22;s__ | 1.25 | 3207293 | 68.75 | 63 | 99.14 | 0.31 |
| p__Bacteroidota;c__Bacteroidia;o__Chitinophagales;f__Chitinophagaceae;g__Niabella;s__ | 1.23 | 4281306 | 38.36 | 390 | 96.15 | 4.62 |
| p__Verrucomicrobiota;c__Kiritimatiellae;o__LD1-PB3;f__CAIVKH01;g__;s__ | 1.20 | 3044093 | 59.81 | 127 | 96.55 | 2.59 |
| p__Proteobacteria;c__Gammaproteobacteria;o__Burkholderiales;f__UKL13-2;g__FEB-7;s__ | 1.19 | 3402912 | 71.55 | 796 | 84.75 | 4.90 |
| p__Armatimonadota;c__Fimbriimonadia;o__Fimbriimonadales;f__Fimbriimonadaceae;g__UBA2387;s__ | 1.07 | 2949814 | 63.1 | 150 | 81.74 | 1.72 |
| p__Proteobacteria;c__Alphaproteobacteria;o__Rhizobiales;f__Rhizobiaceae;g__Aquamicrobium_A;s__ | 0.96 | 3098499 | 68.05 | 209 | 84.51 | 9.05 |
| p__Bacteroidota;c__Bacteroidia;o__Cytophagales;f__Cyclobacteriaceae;g__ELB16-189;s__ELB16-189 sp013414665 | 0.92 | 3861301 | 43.73 | 252 | 93.32 | 6.56 |
| p__Bacteroidota;c__Bacteroidia;o__Chitinophagales;f__Chitinophagaceae;g__Palsa-955;s__ | 0.90 | 3336462 | 40.69 | 26 | 97.44 | 0.00 |
| p__Bacteroidota;c__Bacteroidia;o__Chitinophagales;f__Chitinophagaceae;g__OLB11;s__ | 0.80 | 2420908 | 31.21 | 44 | 98.46 | 0.00 |
| p__Chloroflexota;c__Chloroflexia;o__Thermomicrobiales;f__CADCWN01;g__CADCWN01;s__ | 0.73 | 4504807 | 58.38 | 441 | 97.41 | 0.00 |
| p__Proteobacteria;c__Gammaproteobacteria;o__Burkholderiales;f__Burkholderiaceae;g__Rubrivivax;s__ | 0.53 | 3608138 | 71.23 | 745 | 82.43 | 8.62 |
| p__Chloroflexota;c__Chloroflexia;o__Thermomicrobiales;f__CFX8;g__;s__ | 0.43 | 2754984 | 66.66 | 89 | 92.24 | 1.72 |
| p__Bacteroidota;c__Bacteroidia;o__Chitinophagales;f__Chitinophagaceae;g__Niabella;s__ | 0.37 | 3808670 | 41.35 | 314 | 95.9 | 1.79 |

| **Taxonomy from GTDB (Genome Taxonomy Database)** [*Bacteria* domain] | **Enrich**[1] ($D_z$) | **Length** (bp) | **GC** (%) | **Contigs** | **Comp**[2] (%) | **Cont**[3] (%) |
|---|---|---|---|---|---|---|
| p__Bacteroidota;c__Bacteroidia;o__Chitinophagales;f__UBA10324;g__;s__ | 0.32 | 2683657 | 40.94 | 187 | 92.09 | 3.08 |
| p__Chloroflexota;c__Anaerolineae;o__UCB3;f__;g__;s__ | -0.32 | 6028238 | 66.03 | 1485 | 90.94 | 8.78 |
| p__Bacteroidota;c__Bacteroidia;o__Chitinophagales;f__Chitinophagaceae;g__NS-102;s__ | -0.51 | 4339016 | 39.38 | 105 | 92.26 | 0.10 |
| p__Actinobacteriota;c__Actinomycetia;o__Nanopelagicales;f__UBA10799;g__;s__ | -0.70 | 2148049 | 58.69 | 3 | 99.16 | 0.00 |
| p__Bacteroidota;c__Bacteroidia;o__Chitinophagales;f__LD1;g__UBA7692;s__ | -0.76 | 3124927 | 40.51 | 197 | 98.97 | 3.01 |
| p__Nitrospirota;c__Nitrospiria;o__Nitrospirales;f__Nitrospiraceae;g__Nitrospira_A;s__Nitrospira_A sp001567445 | -0.86 | 3922037 | 60.13 | 163 | 89.94 | 7.14 |
| p__Bacteroidota;c__Bacteroidia;o__Chitinophagales;f__Chitinophagaceae;g__Flavihumibacter;s__ | -0.90 | 3547591 | 52.94 | 825 | 85.26 | 5.34 |
| p__Bacteroidota;c__Kapabacteria;o__Kapabacteriales;f__NICIL-2;g__;s__ | -0.91 | 3577038 | 50.84 | 85 | 100 | 0.00 |
| p__Bacteroidota;c__Bacteroidia;o__Chitinophagales;f__Chitinophagaceae;g__;s__ | -0.98 | 4318938 | 46.23 | 71 | 93.85 | 0.00 |
| p__Bacteroidota;c__Ignavibacteria;o__SJA-28;f__B-1AR;g__UBA2330;s__ | -1.04 | 2978028 | 39.17 | 117 | 96.55 | 0.00 |
| p__Bacteroidota;c__Kapabacteria;o__Kapabacteriales;f__UBA961;g__;s__ | -1.07 | 3110182 | 40.5 | 217 | 98.28 | 0.00 |
| p__Bacteroidota;c__Bacteroidia;o__Chitinophagales;f__Chitinophagaceae;g__JJ008;s__ | -1.09 | 2803149 | 36.27 | 116 | 94.87 | 0.77 |
| p__Proteobacteria;c__Gammaproteobacteria;o__Burkholderiales;f__Rhodocyclaceae;g__Thauera;s__Thauera aminoaromatica | -1.12 | 3863308 | 69.41 | 634 | 80.72 | 5.95 |
| p__Bacteroidota;c__Bacteroidia;o__Chitinophagales;f__Chitinophagaceae;g__Niabella;s__ | -1.15 | 4088796 | 44.23 | 756 | 94.36 | 8.37 |
| p__Chloroflexota;c__Anaerolineae;o__UCB3;f__;g__;s__ | -1.18 | 7188242 | 64.43 | 386 | 91.32 | 0.00 |
| p__Bacteroidota;c__Bacteroidia;o__Chitinophagales;f__Chitinophagaceae;g__NS-102;s__ | -1.18 | 3824646 | 43.08 | 265 | 84.36 | 0.53 |
| p__Bacteroidota;c__Bacteroidia;o__Chitinophagales;f__Chitinophagaceae;g__UTBCD1;s__ | -1.22 | 3819516 | 38.6 | 115 | 98.46 | 1.03 |
| p__Bacteroidota;c__Bacteroidia;o__Chitinophagales;f__Chitinophagaceae;g__UTBCD1;s__ | -1.25 | 3775566 | 38.7 | 114 | 96.92 | 1.03 |
| p__Bacteroidota;c__Bacteroidia;o__Cytophagales;f__Cyclobacteriaceae;g__ELB16-189;s__ | -1.26 | 3054405 | 42.86 | 37 | 83.59 | 1.54 |
| p__Eremiobacterota;c__Xenobia;o__Xenobiales;f__;g__;s__ | -1.34 | 7561854 | 66.76 | 247 | 93.1 | 0.00 |
| p__Bacteroidota;c__Bacteroidia;o__Chitinophagales;f__Chitinophagaceae;g__Ferruginibacter;s__ | -1.35 | 3550783 | 37.14 | 165 | 100 | 0.51 |
| p__Bacteroidota;c__Bacteroidia;o__Chitinophagales;f__Chitinophagaceae;g__UBA1930;s__ | -1.37 | 2542222 | 31.71 | 85 | 93.67 | 4.36 |
| p__Bacteroidota;c__Bacteroidia;o__Chitinophagales;f__UBA10324;g__BJGO01;s__ | -1.38 | 2547451 | 38.92 | 87 | 96.84 | 0.05 |
| p__Bacteroidota;c__Bacteroidia;o__Chitinophagales;f__Chitinophagaceae;g__UBA1930;s__ | -1.40 | 3039471 | 31.54 | 225 | 92.97 | 7.53 |
| p__Myxococcota;c__Polyangia;o__Polyangiales;f__Ga0077539;g__;s__ | -1.41 | 8460108 | 73.89 | 1898 | 80.06 | 6.57 |
| p__Nitrospirota;c__Nitrospiria;o__Nitrospirales;f__Nitrospiraceae;g__Nitrospira_A;s__Nitrospira_A sp009594855 | -1.45 | 3832612 | 58.98 | 261 | 86.21 | 9.49 |
| p__Firmicutes;c__Bacilli;o__Lactobacillales;f__Enterococcaceae;g__Enterococcus_B;s__Enterococcus_B lactis | -1.54 | 3222215 | 38.15 | 217 | 99.61 | 2.62 |
| p__Proteobacteria;c__Gammaproteobacteria;o__Burkholderiales;f__Casimicrobiaceae;g__Casimicrobium;s__ | -1.55 | 3038875 | 66.17 | 368 | 87.85 | 5.77 |
| p__Bacteroidota;c__Bacteroidia;o__Chitinophagales;f__Chitinophagaceae;g__Parafilimonas;s__ | -1.57 | 3049181 | 40.57 | 107 | 99.23 | 6.15 |
| p__Bacteroidota;c__Bacteroidia;o__Cytophagales;f__Cyclobacteriaceae;g__ELB16-189;s__ | -1.94 | 3611179 | 44.13 | 96 | 96.41 | 0.89 |

| **Taxonomy from GTDB (Genome Taxonomy Database)** [*Bacteria* domain] | **Enrich**[1] ($D_z$) | **Length** (bp) | **GC** (%) | **Contigs** | **Comp**[2] (%) | **Cont**[3] (%) |
|---|---|---|---|---|---|---|
| p__Bacteroidota;c__Bacteroidia;o__AKYH767;f__B-17BO;g__;s__ | -1.95 | 3868962 | 33.16 | 366 | 96.41 | 7.69 |

**Notes:**

1. $D_z$ value indicating differential enrichment in treatment versus control reactors. Greater $D_z$ value indicates greater enrichment. $D_z \geq 1.4$ indicates differential enrichment per definition in Section 2.9.3.
2. Completeness (> 80% for medium quality MAGs)
3. Contamination (< 10% for medium quality MAGs)
